# Coordinated changes in gene expression, H1 variant distribution and genome 3D conformation in response to H1 depletion

**DOI:** 10.1101/2021.02.12.429879

**Authors:** Núria Serna-Pujol, Mónica Salinas-Pena, Francesca Mugianesi, François Le Dily, Marc A. Marti-Renom, Albert Jordan

## Abstract

Up to seven members of the histone H1 family may contribute to chromatin compaction and its regulation in human somatic cells. In breast cancer cells, knock-down of multiple H1 variants deregulates many genes, promotes the appearance of genome-wide accessibility sites and triggers an interferon response via activation of heterochromatic repeats. However, how these changes in the expression profile relate to the re-distribution of H1 variants as well as to genome conformational changes have not been yet studied. Here, we combined ChIP-seq of five endogenous H1 variants with Chromosome Conformation Capture analysis in wild-type and H1 knock-down T47D cells. The results indicate that H1 variants coexist in the genome in two large groups depending on the local DNA GC content and that their distribution is robust with respect to multiple H1 depletion. Despite the small changes in H1 variants distribution, knock-down of H1 translated into more isolated but de-compacted chromatin structures at the scale of Topologically Associating Domains or TADs. Such changes in TAD structure correlated with a coordinated gene expression response of their resident genes. This is the first report describing simultaneous profiling of five endogenous H1 variants within a cell line and giving functional evidence of genome topology alterations upon H1 depletion in human cancer cells.

## Introduction

DNA is packaged within the nucleus to efficiently regulate nuclear processes. Chromatin packing involves several hierarchical levels of organization that have been mostly described by chromosome conformation capture techniques, among others. First, at megabases scale, the genome can be segregated into the so-called A and B compartments. The A compartment represents active, accessible chromatin with a tendency to occupy a more central position in the nucleus. The B compartment corresponds to heterochromatin and gene deserts enriched at the nuclear periphery^1^. Second, topological associating domains (TADs), which are submegabase structures, interact more frequently within themselves than with the rest of the genome^2–4^. TADs are conserved across species and cell types and show a coordinated transcriptional status^5,6^. Third, these domains are formed by assemblies of chromatin loops with physical properties that, ultimately, depend on the histone composition and modifications of its resident nucleosomes. In particular, histone H1, which has classically been regarded as a simple condenser, is now known to contribute to the higher-order organization of the genome^7–9^. Histone H1 family is evolutionary diverse and human somatic cells may contain up to seven H1 variants (H1.1 to H1.5, H1.0 and H1X). H1.1-H1.5 variants are expressed in a replication-dependent manner while H1.0 and H1X are replication-independent. H1.2 to H1.5 and H1X are ubiquitously expressed, while H1.1 is restricted to certain tissues and H1.0 accumulates in terminally differentiated cells8,10,11.

Several studies support the idea that H1 variants are not redundant and that functional specificity may exist with H1 variants non-randomly distributed in the genome and interacting with different protein partners^12–18^. For example, in breast cancer cells, knock-down (KD) of each individual H1 variant deregulates different subsets of genes^17,19^. In mouse embryonic stem cells (ESCs), H1c and H1d (orthologs of the human H1.2 and H1.3, respectively) are depleted from high GC/gene-rich regions and are enriched at major satellites^14^. In IMR90 cells, H1.2-H1.5, in contrast to H1.1, are depleted from CpG-dense and regulatory regions^15^, with H1.5 binding correlating with depletion of RNA polymerase II (RNApol II) and repression of target genes in differentiated cells^13^. In skin fibroblasts, H1.0 distribution correlates with GC content and is abundant at gene-rich chromosomes^18^. In T47D breast cancer cells, all H1 variants are depleted at promoters of active genes^16^ and tagged-H1s are enriched at high GC regions with endogenous H1.2 and H1X resulting in opposite profiles. That is, while H1.2 is found in low GC regions and lamina-associated domains (LADs), H1X strongly correlates with GC content and is associated to RNApol II binding sites^16,17^. Moreover, H1.2 and H1X have an opposite distribution among Giemsa bands (G bands), being H1.2 and H1X associated with low and high GC bands, respectively^20^. Finally, a strong correlation has been observed between high H1.2/H1X ratio and the so-called genome B compartment, low GC bands and compact, late-replicating chromatin^20^. Although no functional Hi-C experiments have been performed in H1-depleted human cells, the direct involvement of linker histones in chromatin structure has been proved in mouse ESCs. Hi-C experiments were performed in wild-type and H1-triple knockout (TKO) ESCs. In H1 TKO, an increase in inter-TAD interactions correlated with changes in active histone marks, increased number of DNA hypersensitivity sites and decreased DNA methylation ^21^. These results point to an essential role of histone H1 in modulating local chromatin organization and chromatin 3D organization.

To study the consequences H1 depletion in human cells, we have previously generated a derivative T47D cell line containing a short-hairpin-RNA that affects the expression of several H1 genes as well as the protein levels of mainly H1.2 and H1.4 ^22^. In such cell line, the H1 total levels are reduced to ≈70%, which results in heterochromatic repeats including satellites and endogenous retroviruses overexpression that triggers a strong interferon response. Using this system, here we aim at studying the effects of H1 variant depletion in chromatin organization and nuclear homeostasis. To address this question, we have performed ChIP-seq in T47D breast cancer cells, and Hi-C experiments under basal conditions and after combined depletion of H1.2 and H1.4 (H1 KD). Profiling of endogenous H1 variants revealed that H1.2, H1.5 and H1.0 were abundant at low GC regions while H1.4 and H1X preferentially co-localized at high GC regions. Profiling of H1s within chromatin states showed that all H1 variants were enriched at heterochromatin and low-activity chromatin, but H1X was more abundant at promoters compared to other H1 variants. Finally, H1.4 profile overlapped with H3K9me3 distribution. After H1 KD, chromatin accessibility increased genome-wide, especially at the A compartment where H3K9me3 abundance was reduced. Similarly, the B compartment, where H1.2 was enriched at basal conditions, also showed a more open state. Interestingly, these changes occurred with only slight H1 variant redistributions across the genome. For example, H1.4 profile switched towards the H1.2 group and H1X decreased at heterochromatin and increased in almost all other chromatin states. Our Hi-C results also indicate that upon H1 KD, parts of the genome suffered changes in compartmentalization with no specific direction and TADs increased their internal interactions, which resulted in an increased TAD border strength. In particular, those regions of the genome with high H1.2 overlap resulted in increased local interactions upon H1 KD. Such structural changes were parallel to coordinated gene expression changes within TADs with up-regulated genes enriched in TADs with low basal gene expression and high H1.2 content. Finally, the three-dimensional (3D) modeling of TADs with coordinated gene response indicate that they suffered a general decompaction upon H1 KD. This is the first report describing simultaneous profiling of five endogenous H1 variants within a cell line and giving functional evidence of genome topology alterations upon H1 KD in human cancer cells.

## MATERIALS AND METHODS

### Cell lines, culturing conditions and H1 knock-down

Breast cancer T47D-MTVL derivative cell lines, which carry one stably integrated copy of luciferase reporter gene driven by the MMTV promoter, were grown at 37°C with 5% CO2 in RPMI 1640 medium, supplemented with 10% FBS, 2 mM L-glutamine, 100 U/ml penicillin, and 100 μg/ml streptomycin, as described previously^19^. HeLa and HCT-116 cell lines were grown at 37°C with 5% CO2 in DMEM GlutaMax medium, supplemented with 10% FBS and 1% penicillin/streptomycin. The T47D-MTVL multiH1sh cell line^22^ was used as a model for H1 depletion. This cell line contains a drug-inducible RNA interference system that leads to the combined depletion of H1.2 and H1.4 variants at protein level although it reduces the expression of several H1 transcripts. Construction, establishment and validation of single-H1 knock-downs have been previously described^19^. Specifically, shRNA expression was induced with 6-days treatment of Doxycycline (Dox), in which cells were passaged on day 3. Dox (Sigma) was added at 2.5μg/ml.

### Immunoblot

Chromatin samples were exposed to SDS-PAGE (14%), transferred to a PVDF membrane, blocked with Odyssey blocking buffer (LI-COR Biosciences) for 1 h, and incubated with primary antibodies overnight at 4ºC as well as with secondary antibodies conjugated to fluorescence (IRDye 680 goat anti-rabbit IgG, Li-Cor) for 1 h at room temperature. Bands were visualized in an Odyssey Infrared Imaging System (Li-Cor). Coomassie staining or histone H3 immunoblotting were used as loading controls.

### RNA extraction and reverse transcriptase (RT)-qPCR

Total RNA was extracted using the High Pure RNA Isolation Kit (Roche). Then, cDNA was generated from 100 ng of RNA using the Superscript First Strand Synthesis System (Invitrogen). Gene products were analyzed by qPCR, using SYBR Green Master Mix (Invitrogen) and specific oligonucleotides in a Roche 480 Light Cycler machine. Each value was corrected by human GAPDH and represented as relative units. Specific qPCR oligonucleotide sequences used as previously described^22^.

### Chromatin immunoprecipitation (ChIP)

Chromatin immunoprecipitation was performed according to the Upstate (Millipore) standard protocol. Briefly, cells were fixed using 1% formaldehyde for 10 min at 37ºC, chromatin was extracted and sonicated to generate fragments between 200 and 500 bp. Next, 30 μg of sheared chromatin was immunoprecipitated overnight with the indicated antibody. Immunocomplexes were recovered using 20 μl of protein A magnetic beads, washed and eluted. Cross-linking was reversed at 65ºC overnight and immunoprecipitated DNA was recovered using the IPure Kit (Diagenode). Genomic regions of interest were identified by real-time PCR (qPCR) using SYBR Green Master Mix (Invitrogen) and specific oligonucleotides in a Roche 480 Light Cycler machine. Each value was corrected by the corresponding input chromatin sample. Oligonucleotide sequences are detailed in previous studies^17^.

### ChIP-Seq

#### Library construction and sequencing

Qualified ChIP and Input samples were subjected to end-repair and then 3’ adenylated. Adaptors were ligated to the ends of these 3’ adenylated fragments. Fragments were PCR-amplified and PCR products were purified and selected with the Agencourt AMPure XP-Medium kit. The double stranded PCR products were heat denatured and circularized by the splint oligo sequence. The single strand circle DNA (ssCir DNA) were formatted as the final library and then quality-checked. The library was amplified to make DNA nanoball (DNB) which had more than 300 copies of one molecular. The DNBs were loaded into the patterned nanoarray and single end 50 bases reads were generated in the way of sequenced by combinatorial Probe-Anchor Synthesis (cPAS).

#### ChIP-seq data analysis

Single-end reads were quality-checked via FastQC (v0.11.9) and aligned to the human GRCh37/hg19 reference genome using Bowtie2 (v2.3.5.1)^23^ with default options. SAMtools (v1.9)^24^ utilities were used to filter out the low-quality reads with the flag 3844. Input, H1 variants, and H3K9me3 genome coverage was calculated and normalized by reads per million with BEDTools (v2.28.0)^25^, and regions with zero coverage were also reported in the ChIP-Seq annotation (*genomecov -ibam -bga -scale*). MACS2 (v2.1.2)^26^ was used to subtract input coverage from H1 variants and H3K9me3 to generate signal tracks (*bdgcmp -m subtract*). In the case of H3K9me3 samples, peaks were also called with MACS2 (*callpeak --broad --nomodel --extsize*). SICER (v1.1)^27^ was used to identify histone ChIP-enriched regions with the following parameters: *redundancy thereshold* = 1, *window size* = 200, *fragment size* = 150, *effective genome fraction* = 0.80, *gap size* = 200 and *FDR* = 0.01. Genomic annotation of the identified H1-enriched regions was performed with CEAS software^28^.

ChIP-Seq data on histone H1 variants and H3K9me3 epigenetic modification from T47D multiH1sh cells treated or not with Dox has been deposited in NCBI’s Gene Expression Omnibus and is accessible through GEO Series accession number GSE156036. ChIP-Seq data on histone H1 variants from WT T47D cells is at GSE166645.

### Antibodies

Specific antibodies recognizing human H1 variants used for ChIP/ChIP-seq were: anti-H1.0/H5 clone 3H9 (Millipore, 05-629-I), anti-H1.2 (Abcam, ab4086), anti-H1.4 (Invitrogen, 702876), anti-H1.5 (Invitrogen, 711912) and anti-H1X (Abcam, ab31972). ChIP-seq of H3K9me3 was performed using anti-H3K9me3 (Abcam, ab8898). Other antibodies used were: anti-H1.0 (Abcam, ab11079), anti-H1.3 (Abcam, ab24174), anti-H1.5 (Abcam, ab24175) and anti-H3 (Abcam, ab1791).

### In situ Hi-C

Hi-C libraries were generated from T47D multiH1sh cells treated or not with Dox, as single replica (r1) or duplicate (r2 and r3), as previously described ^29,30^. In brief, adherent cells were cross-linked with 1% formaldehyde in PBS for 10 min at room temperature and glycine 0.125 M was added for 5 min at room temperature and for 15 min at 4ºC to stop the cross-link reaction. Before permeabilization, cells were treated for 5 min with trypsin. Nuclei digestion was performed with 400 units of MboI restriction enzyme. The ends of restriction fragments were labeled using biotinylated nucleotides and ligated with T4 DNA ligase. After reversal of cross-links, DNA was purified and sheared (Diagenode BioruptorPico) to obtain DNA fragments between 300 and 500 bp and ligation junctions were pull-down with streptavidin beads. Hi-C libraries were amplified, controlled for quality and sequenced on an Illumina HiSeq 2500 sequencer (r1) or DNBseq (r2,r3).

### Hi-C data pre-processing, normalization and generation of interaction matrices

The analysis of Hi-C data, from FASTQ files mapping to genome segmentation into A/B compartments and TADs, was performed using *TADbit*^31^, which started by performing a quality control on the raw data in FASTQ format. Next, sequencing reads were mapped to the reference genome (GRCh37/hg19) applying an iterative strategy and using the GEM mapper^33^. Mapped reads were filtered to remove those resulting from unspecified ligations, errors or experimental artefacts. Specifically, nine different filters were applied using the default parameters in TADbit: self-circles, dangling ends, errors, extra dangling-ends, over-represented, too short, too long, duplicated and random breaks^31^. Hi-C data were next normalized with OneD correction to remove Hi-C biases and artifacts^32^. Filtered read-pairs were binned at the resolutions of 1 Mb, 500 kb, 100 kb and 10 kb, applying biases from the normalization step and decay correction to generate interaction matrices. Hi-C data on T47D breast cancer cells has been deposited in NCBI’s Gene Expression Omnibus and is accessible through accession number GSE172618. A summary of the number of valid reads obtained per replica and filtered artifacts is shown as **Suppl. Table 1**. Replicates were compared and merged with *TADbit merge* that implements the HiCRep score^34^.

### Genome segmentation into Topologically Associating Domains (TADs)

TADs were identified at the resolution of 50kb using *TADbit segment* with default parameters. Briefly, TADbit segments the genome into constitutive TADs after analyzing the contact distribution along the genome using a BIC-penalized breakpoint detection algorithm^31^. This algorithm leads to a ∼99% average genome coverage. To assign a strength value to each TAD border, TADbit repeats the dynamic programming segmentation 10 times after the optimum is reached, each time decreasing the by a fix off-set the optimal TAD border detection path. The strength of a TAD border is then calculated as the number of times it was included in the optimal pathway. If a TAD border is found in all 10 sub-optimal paths, then the score of the border is equal to 10, if it was found only one time, the score is 1. Finally, TADbit also returns a TAD density score as the ratio between the number of interactions within TADs and the number of interactions of the rest of the genome.

### Genome segmentation into A/B compartments

A/B compartments were identified at 100kb resolution using HOMER^35^. Briefly, HOMER calculates correlation between the contact profiles of each bin against each other, and performs principal component analysis (PCA) on chromosome-wide matrices. Normally, the A compartment is assigned to genomic bins with positive first principal component (PC1), and the B compartment is assigned to genomic bins with negative PC1. However, in some chromosomes and in cell lines with aberrant karyotypes, the PC1 is reversed in the sign, with A compartment corresponding to negative PC1, and B compartment corresponding to positive PC1. Additionally, sometimes the PC1 captures other correlations in the chromosome that do not correspond to the compartments. For these reasons, all PC1 and PC2 for all chromosomes were visually inspected and correctly assigned to decipher the proper segmentation of the genome into the A and B compartments.

### 3D modelling of TADs based on Hi-C data

*TADbit model*^31^ was used with default parameters to generate 3D models of selected TADs at the resolution of 10 kb. Hi-C interaction maps were transformed into a set of spatial restraints that were then used to build 3D models of the TADs that satisfied as best as possible the imposed restraints, as previously described^36,37^. For each TAD, we generated 1,000 models, structurally aligned and clustered them in an unsupervised manner, to generate sets of structurally related models. For every TAD, we used the main cluster to compute consistency, accessibility, density, radius of gyration, and walking angle^31^. Consistency quantifies the variability of the position of particles across the considered set of models. Accessibility measures with a fraction from 0 to 1 how much each particle in a model is accessible to an object (*i*.*e*. a protein complex) with a radius of 100 nm. Density measures a proxy for local DNA compactness as the ratio of DNA base pairs and the distances between two consecutive particles in the models – the higher the density, the more compact the DNA. Walking angle measures the angle between triplets of consecutive particles —the higher the value, the straighter the models— and can be used as a proxy for the stiffness of the chromatin fiber. Finally, radius of gyration measures 3D structure compactness as the root mean square distance of the all particles in a model from its center of mass.

### ATAC-Seq data analysis

ATAC-Seq data identified by the accession number GSE100762 was reprocessed as previously described ^38^ with slight modifications. Paired-end reads were quality-checked via FASTQC (v0.11.9), trimmed, and subsequently aligned to the human GRCh37/hg19 reference genome using Bowtie2 (v2.3.5.1) ^23^. SAMtools (v1.9)^24^ was used to filter out the low-quality reads with the flag 1796, remove reads mapped in the mitochondrial chromosome and discard those with a MAPQ score below 30. The peak calling was performed with MACS2 (v.2.1.2)^26^ by specifying the -*BAMPE* mode. Filtered BAM files were also used to compute the ATAC-Seq genome coverage, which was normalized to reads per million (*genomecov -ibam -bga -scale*).

### Genomic data retrieval

Genome-wide GC content, G bands coordinates at 850 bands per haploid sequence (bphs) resolution and chromosomes coordinates were obtained from the UCSC human genome database^39,40^. G bands were classified as G positive (Gpos25 to Gpos100, according to its intensity upon Giemsa staining), and G negative (unstained), which were further divided into four groups according to their GC content (Gneg1 to Gneg4, from high to low GC content) (**Suppl. Figure 2A**). HeLa-S3 genome segmentation by ChromHMM (ENCODE) was obtained from UCSC human genome database^39,40^. RNA-seq and ATAC-seq datasets were download from GEO (accession numbers GSE83277 and GSE100762, respectively) an parsed as previously described^22^.

## Results

### A stable genome distribution of H1 variants correlates with GC content and chromatin state

It has been previously described that the content of histone H1 variants varies between cell types and along differentiation^8,41^. Moreover, its genomic distribution is non-homogeneous and with specific patterns depending on the variants^13–18,20^. Therefore, we hypothesize that altering the H1 variants composition in a particular cell type may affect the genomic distribution of the different variants. To test this, we performed ChIP-seq experiments in T47D cells harboring an inducible multiH1 shRNA expression vector which, upon Doxycycline treatment, efficiently depletes H1.2 and H1.4 proteins (H1 KD)^22^. After testing the efficacy of H1 KD by Western blot (**Figure 1A**), we performed ChIP with antibodies against endogenous H1.2, H1.4, H1.5, H1.0 and H1X. The amount of DNA immunoprecipitated with H1.2 and H1.4 antibodies decreased >65% in treated cells compared to untreated, confirming the antibody specificity and the effect of the H1 knock-down. ChIPed DNA was qPCR-amplified with oligonucleotides for TSS and distal promoter regions of *CDK2* (active) and *NANOG* (inactive) genes (**Figure 1B**), which confirmed that all ChIPs efficiently worked compared to unspecific IgG. The active gene presented the characteristic H1 valley at the Transcription Starting Site (TSS) compared to the distal region, but not the inactive gene^16^. Upon H1 KD, the signal of H1.2 and H1.4 significantly decreased, while H1.0 signal increased, in agreement with the Western blot results (Figure 1A-B). The effect of H1 KD as well as the specificity of H1 antibodies were further confirmed by Western blot, ChIP-qPCR and RT-qPCR (**Suppl. Figure 1**).

**Figure 1.**
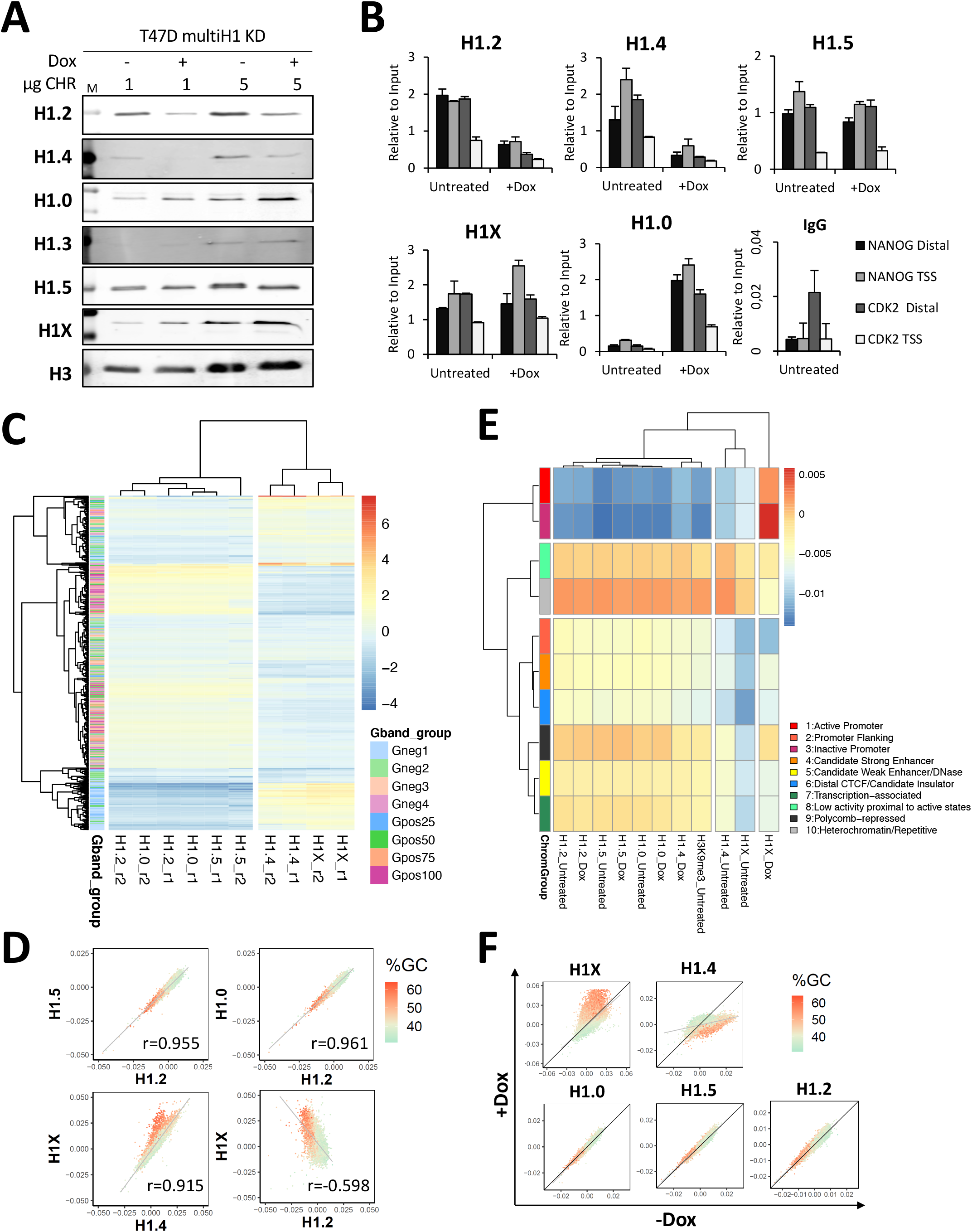
Genomic distribution of histone H1 variants upon H1 knock-down in breast cancer cells. **(A)** Immunoblot analysis of H1 depletion in multiH1 KD cells. Chromatin extracts (1 or 5µg of protein) from T47D multiH1 KD cells cultured in the presence or not of Doxycycline for 6 days were run in SDS/PAGE and immunoblotted with the indicated antibodies against H1 variants or histone H3 as loading control. **(B)** ChIP-qPCR of H1 variants in multiH1 KD cells. Chromatin from untreated or Dox-treated H1 KD cells was used for ChIP with antibodies against H1 variants and unrelated IgG as a control. Resulting DNA was amplified by qPCR with oligos for distal promoter (3kb upstream TSS) and TSS regions of genes *CDK2* and *NANOG*. ChIP amplification is shown relative to input DNA amplification. A representative experiment quantified in triplicate is shown. **(C)** Heat map and cluster analysis of the input-subtracted ChIP-seq abundance of H1 variants within Gpos and Gneg bands from untreated T47D cells. Two main clusters of H1 variant distribution are formed, one with abundant H1.X and H1.4 at high GC bands, the other with H1.2, H1.5 and H1.0 enriched at low GC bands. Two replicates are shown (r1, r2). **(D)** Scatter plots of the indicated H1 variant pairs input-subtracted ChIP-seq abundance within 100-kb bins of the human genome. The GC content at each bin is color-coded. Pearson’s correlation coefficient is shown (*P*-value<0.001). **(E)** Heat map and cluster analysis of the input-subtracted ChIP-seq abundance of H1 variants from WT or H1 KD T47D cells (-/+Dox) within 10 chromatin states (ChromHMM segmentation). The profile of H3K9me3 from WT cells is included. **(F)** Scatter plots of H1 variants input-subtracted ChIP-seq abundance within 100-kb bins of the human genome in multiH1 KD cells treated or not with Doxycycline. The GC content at each bin is color-coded.

We have previously shown that H1.2/H1X ratio strongly correlates with B compartment, late replicating, inaccessible chromatin and low GC bands^20^. To further extend this analysis, we measured the abundance of each H1 within G bands and compared its distribution upon H1 KD. Unsupervised clustering of H1 variant distributions in G bands clearly show the existence of two major clusters of H1 variants within G bands (**Figure 1C** and **Suppl. Figure 2A**). In untreated cells, H1.2 was enriched towards low GC content regions (that is, Gpos100/Gneg4, repressed bands), and H1X was enriched at high GC (that is, Gpos25/Gneg1, active bands). Additionally, H1.4 was also enriched at high GC bands, whereas H1.5 and H1.0 were enriched towards low GC bands. These results confirm and expand previous findings on the distribution of H1 variants in the genome^16,20^. Interestingly, correlation analysis of the distribution of H1 variants genome-wide using bins of 100-kb confirmed the existence of these two groups of variants (*i*.*e*., H1.2, H1.5 and H1.0 in low GC regions as well as H1.4 and H1X in high GC regions) with H1.2 and H1X selected as prototypes of the two groups and opposed distribution within the genome (**Figure 1D** and **Suppl. Figure 2B-C**). Next, we assessed whether the clustering distribution of H1 variants would also correlate with genomic chromatin states. To do so, we used as a proxy several genomic datasets for HeLa-S3 cell lines, which have previously been used to generate a 10-chromatin state (colors) map ^42^. Most H1 variants were particularly abundant within *heterochromatin/repetitive* and *low-activity* chromatin states, but also at *polycomb-repressed, transcription-associated* and *weak-enhancer* (**Figure 1E** and **Suppl. Figure 2D**). In concordance, such variants were underrepresented at *active* and *inactive promoter* states. This trend was broken by the H1X variant, which was enriched at *promoters* confirming that this variant is the most specific of all with respect its genome-wide distribution. H1.4 was the variant that overlapped the most with H3K9me3 profile within chromatin states.

Changes of H1 variants distribution upon H1 KD were further analyzed within 100-kb bins throughout the genome. Upon H1 KD, H1.0 distribution was unaltered, while H1.2 and H1.5 were slightly increased specially at high GC bins. H1X occupancy increased at high GC bins and decreased at low GC bins, whereas H1.4 decreased at high GC bins (**Figure 1F** and **Suppl. Figure 2A**). Similarly, upon H1 KD, H1.2, H1.5 and H1.0 profiles within chromatin states were not altered and H1X profile decreased at *heterochromatin* and increased in almost all other chromatin states, particularly at *polycomb-repressed* regions and *promoters*, and among them the highest increase occurred at *inactive promoters* (**Figure 1E** and **Suppl. Figure 2D**). Finally, H1.4 profile switched towards the H1.2 group.

In summary, ChIP-seq data in T47D cells demonstrated that H1 variants are differentially distributed through the genome in two profiles: H1.2, H1.5 and H1.0 enriched towards low GC regions and H1X and H1.4 more abundant at high GC regions. Upon H1.2 and H1.4 depletion, H1.2, H1.0 and H1.5 did not significantly change their genomic distribution, whereas H1X increased at high GC regions, where H1.4 was selectively depleted. H1.0, whose expression and protein levels increased, was homogeneously incorporated throughout the genome.

### Changes on genome architecture upon depletion of multiple histone H1 variants

Chromosome conformation capture techniques such as Hi-C allows to detect local and distal contacts within the genome and to establish the position of borders flanking the so-called topologically associating domains (TADs). Hi-C experiments also allow to establish a division of the genome into two compartments, active (A) and inactive (B). To address the consequences of histone H1 depletion on genome architecture, we prepared nuclear DNA from untreated and 6-days Doxycycline-treated multiH1 KD cells, in two independent experiments with a total of 3 replicates, and performed the Hi-C protocol (**Suppl. Figure 3**). After assessing the similarity between Hi-C replicates using HiCRep score (**Materials and Method**s and **Figure 2A**), replicates within samples were merged and analysed as a single experiment for WT and H1 KD. Analysis of the average Hi-C interactions as a function of genomic distance indicates that upon H1 depletion there was a decrease in short and medium-range interactions (<30 Mb), and an increase in long-range contacts (>30 Mb) (**Figure 2B**). To further characterize where those average changes occurred, we segmented the genome first into compartments and then into TADs for WT and H1 KD samples.

**Figure 2.**
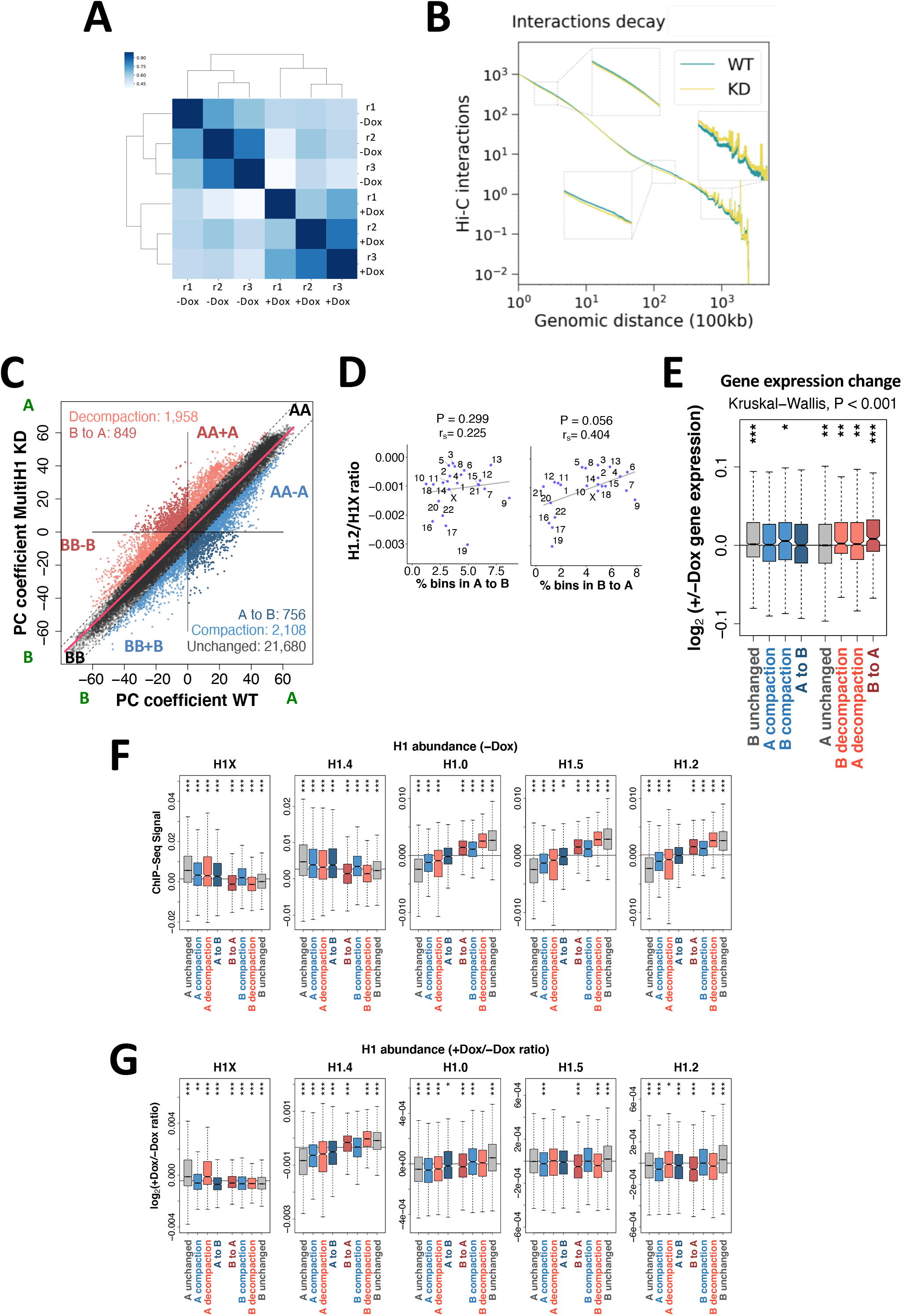
A/B compartments redistribution upon multiH1 KD. **(A)** Hierarchical clustering of Hi-C replicates from WT (-Dox) and multiH1 KD (+Dox) cells based on the Hi-C reproducibility score between paired experiments. **(B)** Plot comparing the distribution of Hi-C interactions versus genomic distance across the genome for a maximum distance of 500 Mb for WT and H1 KD cells. **(C)** Scatter plot of principal component (PC) coefficients for 100-kb genomic segments (bins) from WT (-Dox) and multiH1 KD (+Dox) cells. PC coefficients were used to define A (positive PC) and B (negative PC) compartments, as well as compartment shifting (A-to-B and B-to-A), compaction (blue bins AA-A and BB+B) and decompaction (red bins AA+A and BB-B) upon H1 KD. Unchanged segments AA and BB are black-colored. A polynomial regression line was used to model the relationship between the dependent and the independent variables. **(D)** A/B compartments redistribution within chromosomes. Scatter plot between the percentage of bins that changed from A to B or vice versa upon H1 KD, and the average H1.2/H1X ChIP-seq signal ratio within TADs, for each chromosome. Spearman’s correlation coefficient is shown as well as *P*-value. **(E)** Gene expression changes upon H1 KD within bins changing compartment or compaction rate. Normalized RNA-seq reads of coding and non-coding genes before and after Dox-induced H1 KD within 100-kb bins of the 8 categories obtained in (C) were used to calculate the +/-Dox fold-change (expressed as log2). **(F-G)** Box plot showing H1 variants input-subtracted ChIP-seq signal within bins of each category in WT cells (-Dox) (**F**), or the ratio of change (log2) in H1 KD (+Dox) compared to untreated cells (-Dox) (**G**). (***) P < 0.001; (**) P < 0.01; (*) P < 0.05. Kruskal-Wallis test determined that there were statistically significant differences between the groups (P < 0.001). One-sample Wilcoxon signed-rank test was used to compare each group of bins against the median gene expression changes (E), H1 variants input-subtracted ChIP-seq signal (F), or ratio of change (G).

The segmentation of the genome into the A and B compartments remained largely unchanged upon H1 KD (∼80% of the 100-kb bins did not change compartment, **Figure 2C**). However, significant differences in compartmentalization were observed. For example, about 280Mb of the genome decompacted (B to A direction) after H1 KD with 1/3 of the bins moving from the B compartment to an A compartment. Conversely, about 294Mb of the genome compacted (A to B direction) with about 1/4 completely changing compartment category (**Figure 2C**). Interestingly, these changes in compartmentalization were not homogenous across the genome, being B-to-A shifts upon H1 KD more frequent within chromosomes with high H1.2/H1X ratio. Notably, the expected anti-correlation for bins moving from the A compartment to the B compartment was not observed (**Figure 2D**). To assess if changes in compartment were related to gene activity, we also explored whether gene expression was altered within bins changing compaction upon H1 KD using RNA-seq data previously acquired in the same cell systems^22^. Significant overall gene up-regulation was observed within bins being decompacted (B-to-A and A or B decompaction), but the opposite was not observed for bins being compacted (**Figure 2E**). We next wondered whether the changes in compartmentalization and expression were dependent on the basal distribution of H1 variants in the genome as well as their re-distribution upon H1 KD. As expected, we found that H1X and H1.4 were enriched in the A compartment and H1.0, H1.5 and H1.2 were enriched in the B compartment (**Figure 2F**). Interestingly, such a trend was pronounced for all the bins in the genome which compartmentalization did not change upon H1 KD indicating that the basal state of different H1 variants could determine how compartments respond to H1 depletion. However, was the observed trend upon H1 depletion also accompanied by a change of H1 variant distribution? Interestingly, H1X decreased upon H1 KD in B compartment bins (regardless of their change in compartmentalization) as well as in A compartment bins that compacted or even moved to the B compartment (**Figure 2G**), which could indicate that decrease of H1X is associated to B compartmentalization. Similarly, H1.2 decreased in all A compartment bins as well as B compartments that decompacted or even moved to the A compartment, which again indicates that H1.2 decrease is associated to A compartmentalization. To note that, despite H1.4 clear depletion after H1 KD, its changes associated to compartmentalization did not correlate with the observed changes in H1X (**Figure 2G**). In fact, H1.4 decreased in all A compartment bins and increased in all B compartment bins after H1 depletion, which could indicate a redistribution that could play a significant role in compartmentalization.

Topologically Associating Domains or TADs comprise the next scale of the so-called higher-order organization of chromatin after compartmentalization^43^. Similar to the compartmentalization changes, the large majority of TAD borders (*i*.*e*., 71.0%) remained unchanged upon H1 KD, 12.4% shifted by only one bin, and 16.5% were not conserved (that is, shifted by more than 1 bin, newly formed or disappeared; **Figure 3A** as example of a de novo detected border after H1 KD). To determine whether those changes could be linked to the basal distribution of H1 variants prior H1 KD, we interrogated the TAD enrichment of H1.2 or H1.4, which we identified above having a role in A/B compartmentalization. The results indicate that H1.2 was significantly depleted at non-conserved compared to conserved TAD borders and H1.4 was higher at TADs with non-conserved borders (**Figure 3B**). Interestingly, the differences in border position were also associated to changes in border strength. Upon H1 KD there was an increase in border strength for conserved and shifted TAD borders but not for the non-conserved borders, which slightly decreased its border strengths but with no statistically significant differences (**Figure 3C**). The results suggest, thus, that “soft” borders were prone to be altered upon histone H1 depletion, both in its position as well as in its strength.

**Figure 3.**
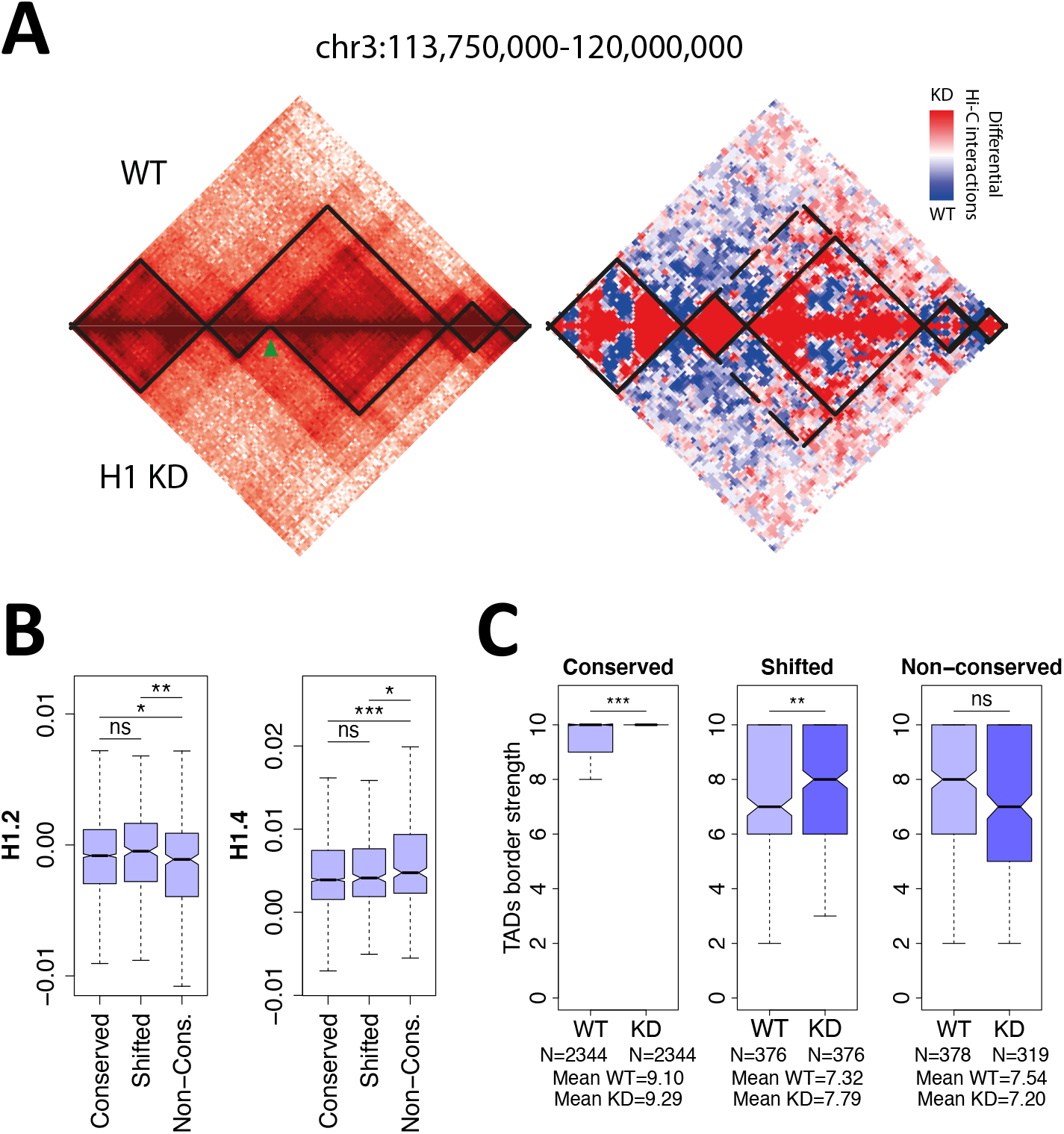
TAD boundaries changes upon H1 KD. **(A)** Hi-C interaction maps of 6.25Mb region in chromosome 3 at 50-kb resolution. *Left panel* is a heat map of Hi-C maps normalized by reads coverage in Log2 scale with TADs overlayed by black lines. Top triangle of the map corresponds to Hi-C in WT and lower triangle to H1 KD. Green arrow points to the de-novo detected TAD border in H1 KD. *Right panel*, differential Hi-C map showing the enrichment of internal interaction in the two separated TADs around the new detected border. **(B)** Box plot showing the H1.2 and H1.4 input-subtracted ChIP-seq signal in WT cells within TADs containing the TAD borders divided in conserved, shifted <100kb and non-conserved according to their behavior upon H1 KD. **(C)** TAD border dynamics. Box plot of normalized border strength distribution for TAD borders in WT and H1 KD cells, divided in conserved, shifted <100kb and non-conserved borders. (***) P < 0.001; (**) P < 0.01; (*) P < 0.05 (Mann-Whitney test).

The observed increased border strength was associated to an increase in intra-TAD (*i*.*e*., local interactions) both within A and B compartments and a decrease of inter-TAD interactions (*i*.*e*., non-local interactions) within the A and between A and B compartments (**Figure 4A**). The increase of local interactions (intra-TAD) with a decrease of non-local interactions (inter-TAD) was also observed with the *D-score*, which measures the differential local interactions per each of the 100-kb bins in a Hi-C matrices. Specifically, the *D-score* is the average of differential interactions between WT and H1 KD of each bin with any other bin within a window of 2 Mb. Thus, it measures if a bin is surrounded by mainly a region in the genome of increased (*D-score*>0) or decreased (*D-score*<0) interactions (**Figure 4B**). Next, we compared the basal distribution of H1 variants with the D-score and found that H1.2 signal was a strong predictor of the *D-score* across the genome (corr.coef.=0.707 and **Figure 4B,C**). Conversely, there was an inverse correlation between the *D-score* and the basal abundancy of H1X variant (corr.coef.=-0.422). In other words, those regions of the genome with high H1.2 overlap are likely to result in increased local interactions once H1 is depleted while regions with high H1X are likely to decrease interactions.

**Figure 4.**
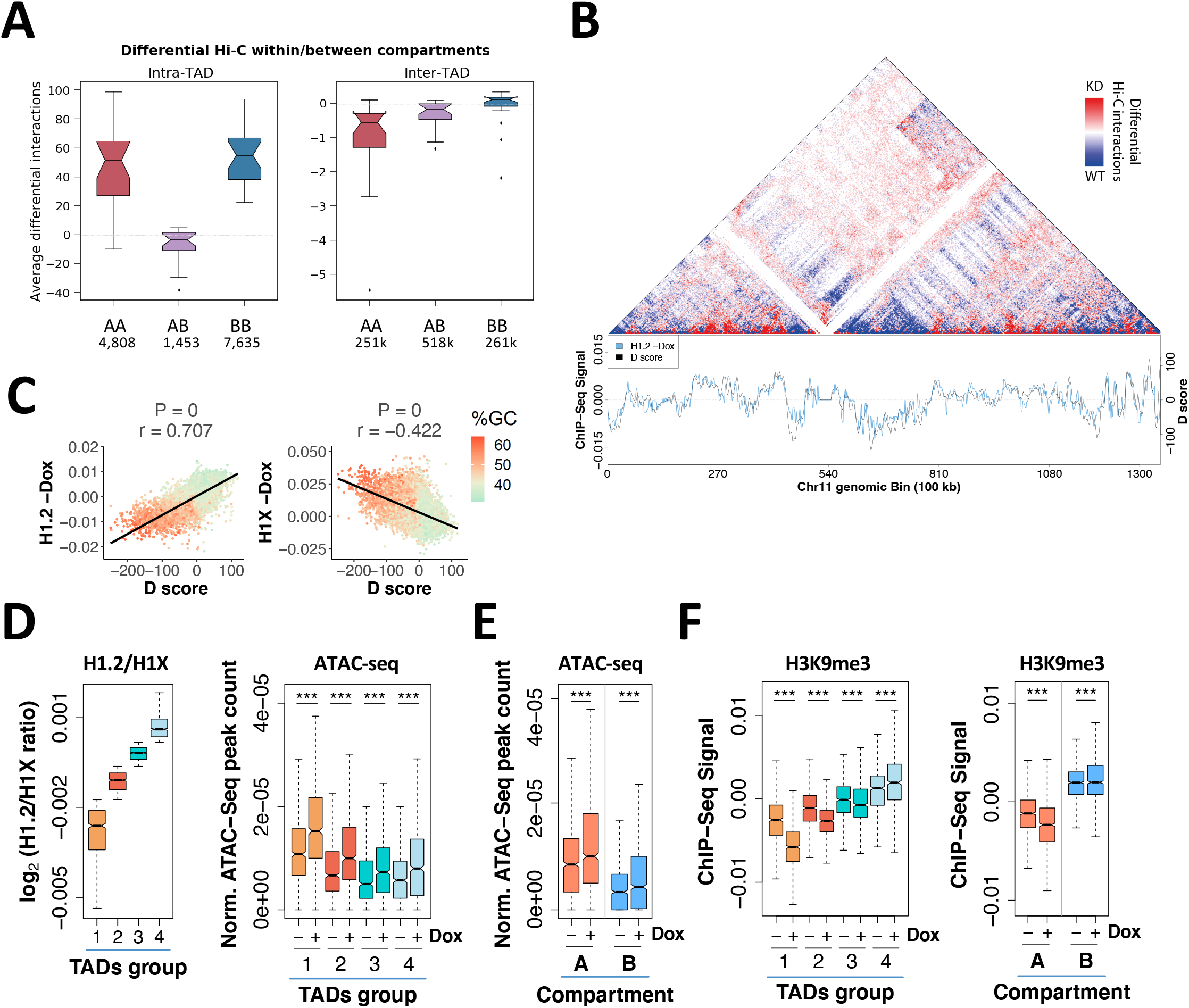
Dynamics of Hi-C genomic interactions and chromatin changes upon H1 KD. **(A)** Box plot showing average number of differential intra-or inter-TAD interactions per chromosome upon H1 KD in different compartments, at 100 kb resolution. The average number per chromosome of differential interactions for each category is indicated in the ticks of X axes. **(B)** *Top*-Differential Hi-C map. Increased (red colored) and decreased (blue colored) interactions in contact matrices of chromosome 11 (0-135Mb) of H1 KD compared to WT cells, at 100kb bins resolution. *Bottom*-*D* score. Profiles of differential interaction *D* score and input-subtracted H1.2 ChIP-seq abundance from WT cells along chromosome 11, calculated within 100 kb bins. **(C)** Scatter plots between differential interaction *D* score and H1.2 or H1X abundance from WT cells, genome-wide. Spearman correlation coefficient is shown as well as *p*-value. The GC content at each bin is color-coded. **(D-E)** Box plots showing the relative number of ATAC-seq peaks (normalized by length) within TADs classified according to H1.2/H1X ratio (Groups 1 to 4) (**D**) or within A/B compartments (**E**), at WT and H1 KD (-/+Dox) cells. The ChIP-seq H1.2/H1X signal ratio within TADs in the 4 groups reported is shown for reference in (D). **(F)** Box plots showing the H3K9me3 input-subtracted ChIP-seq signal within TADs classified according to H1.2/H1X ratio (left) or within A/B compartments (right), at WT and H1 KD (-/+Dox) cells. (***) P < 0.001; Wilcoxon signed-rank test. A compartment, N = 1,057; B compartment, N = 1,009; TADs, N = 735 TADs per group.

Next, to identify if there was a correlation between the observed changes in H1 variants upon H1 KD within the spatial genome and the underlying chromatin state, we further classified TADs by their content in H1.2 and H1X variants (that is, we generate four discreate groups of TADs from lowest to highest H1.2/H1X ratio; **Figure 4D**). Upon H1 depletion, accessibility measured by ATAC-seq was significantly increased at all TAD categories, but its increase was more pronounced at low H1.2/H1X TADs (**Figure 4D**). Accordingly, accessibility was also increased at the A and B compartments but most notably in A compartment (**Figure 4E**). As expected, the opposite trend was observed in analyzing the distribution of the repressive mark H3K9me3 upon H1 KD. Indeed, H3K9me3 ChIP-seq signal decreased more in the A compartment compared to the B compartment and in low H1.2/H1X ratio TADs (**Figure 4F**), which indicates again that chromatin decompaction upon H1 depletion occurs more prominently in already open regions of the genome.

Altogether our findings indicate that the genome structure is not generally but specifically altered upon depletion of H1 variants. First, local and non-local interactions genome-wide were differentially altered with short and mid-range interactions decreasing and long-range increasing. Second, these changes in interactions correlated with changes in A and B compartments associated to changes of gene expression. Third, intra-TAD interactions increased, mostly within A or B compartments, which resulted in a clear increase of TAD border strength. Fourth, these genome interaction changes were more prominent depending on the basal H1 variant occupancy being the distribution of H1.2 and H1X most informative of the observed changes. Fifth, and final, depletion of H1 variants resulted in an overall increase of accessibility of chromatin, which also depended on the basal occupancy of H1.2 and H1X.

### Gene expression is coordinately altered within TADs upon H1 KD

As previously observed, H1 variant depletion resulted in deregulation of hundreds of genes with about one third of the up-regulated genes associated to transcriptional response to interferon^22^. In our experiments, a total of 1,089 and 1,254 genes were up-regulated and down-regulated, respectively (FC ≥ 1.4, adjusted *P*-value ≤ 0.05, **Figure 5A**). Interestingly, groups of regulated genes were more often than expected co-localized within the same TAD. The 2,343 deregulated genes were distributed across 1,292 TADs with an enrichment of TADs with either only up or down regulated genes. For example, there was 531 TADs with at least one up-regulated genes and no down-regulated genes (here called “Up”). Similar numbers were observed for down-regulated TADs with 520 with at least one down-regulated gene and no up-regulated genes (here called “Dw”). Finally, a total of 241 TADs contained at least 2 genes deregulated with mix directions (here called “UpDw”). UpDw TADs corresponded to higher gene density and lower H1.2/H1X ratios compared to either TAD-Up or TAD-Dw. Finally, TADs without deregulated genes (here called Control) had the highest H1.2/H1X ratio as well as the lowest gene richness (**Figure 5B**). Most TADs containing significantly deregulated genes upon H1 KD were located before KD within the A compartment (**Figure 5C**), while Control TADs were enriched at the B compartment. Accordingly, H1X was significantly enriched within UpDw TADs and depleted from Control TADs contrary to the observed trend for H1.2 variant. Chromatin remodeling also followed the expected trends for the TAD groups classified by their change in expression of the resident genes. For example, upon H1 KD, ATAC-seq accessibility increased globally in all TADs, especially in UpDw type (**Figure 5D**). Conversely, H3K9me3 abundance significantly decreased in Dw and UpDw TADs (**Figure 5E**).

**Figure 5.**
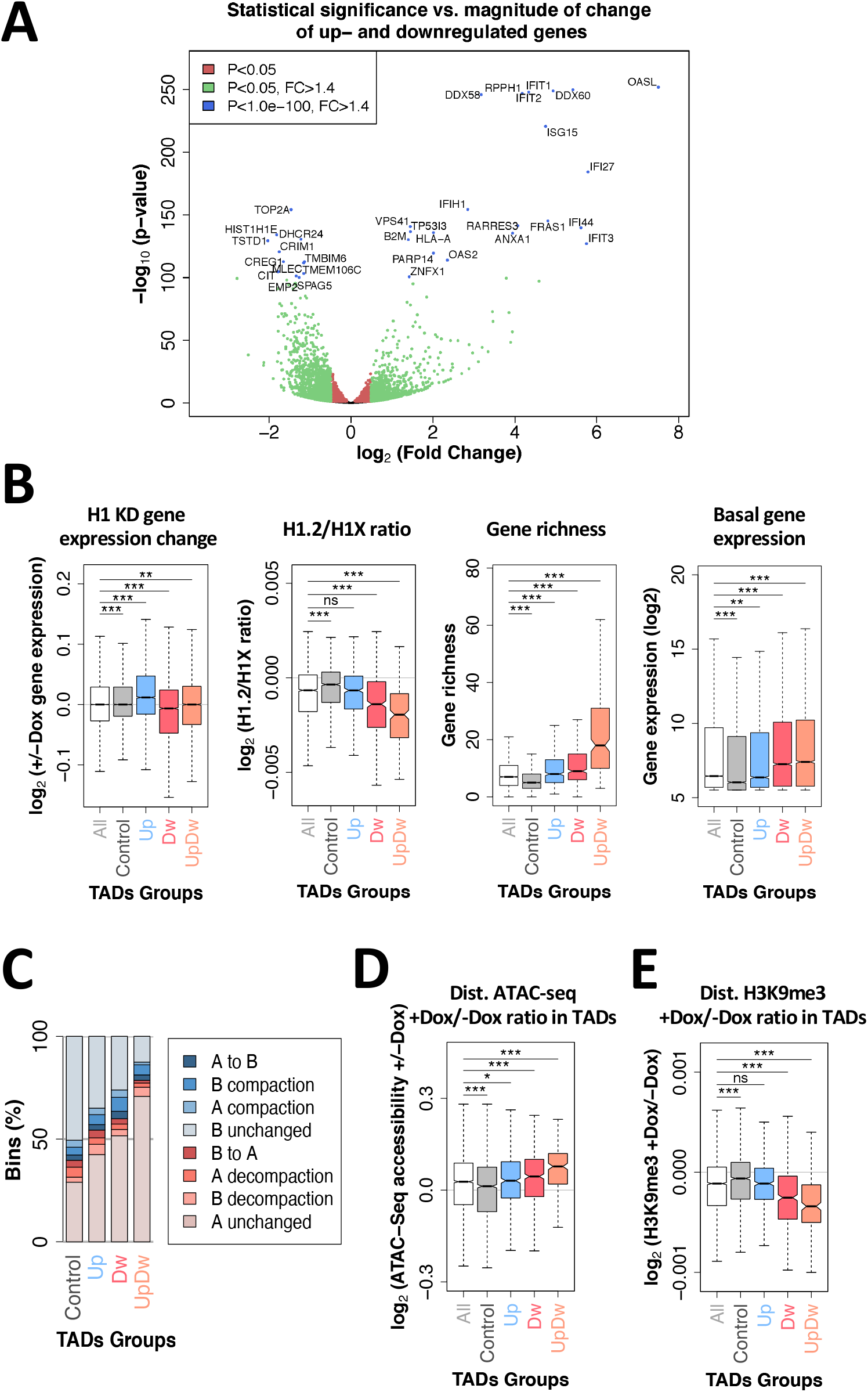
Characterization of TADs containing genes up- or down-regulated upon multiH1 KD. **(A)** Volcano plot of gene expression changes upon multiH1 KD measured by RNA-seq. Log2 of fold change is plotted against the minus log10 of *P*-value. Green dots mark genes with a FC≥±1.4 and *P*<0.05. Interferon stimulated genes (ISGs) (*P*-value<1e-100) are labeled (blue dots). **(B)** Box plot showing the gene expression change upon H1 KD (+/-Dox), Log2 of ChIP-seq H1.2/H1X ratio (data from WT cells), normalized gene richness and normalized basal gene expression within TAD groups: all TADs, TADs without deregulated genes upon H1 KD (Control), TADs containing only up-regulated genes, only down-regulated, or both up- and down-regulated genes simultaneously (FC≥±1.4, *P*<0.05). **(C)** Bar plots showing the frequency of overlap between all the TAD groups described in (B) and genome segments within A/B compartment categories described in Figure 2C. **(D-E)** Box plot showing ATAC-seq accessibility gain (**D**) and changes in H3K9me3 ChIP-seq signal (**E**) upon H1 KD (+/-Dox) within the TAD groups described in (B). (***) P < 0.001; (**) P < 0.01; (*) P < 0.05 (Mann-Whitney test).

As previous described in T47D cell lines^6^, we observed an intra-TAD coordinated response of gene expression. Indeed, we found an enrichment of gene-rich TADs (that is, with at least 4 genes) where most of its genes changed expression in the same direction. Specifically, TADs with over 70% of their genes up-regulated or at least 80% down-regulated were observed in proportions beyond random expectation (**Figure 6A**). These correspond to TADs where all or most of the genes changed expression in the same direction upon H1 KD, including Interferon stimulated genes (ISGs) such as ISG20, CMPK2, DDX60 or GBP3 (**Figure 6B,C** and **Suppl. Figure 4A**). This could result from two hypothetical scenarios: i) upon H1 depletion, the whole TADs were (architecturally) affected and most resident genes became up-or down-regulated coordinately; ii) upon H1 depletion, some gene within a TAD became deregulated and, consequently, neighbor genes within the same TAD changed expression in the same direction. To discern between these two scenarios, we characterized the groups of TADs with most coordinated changes of expression upon depletion of H1. Generally, these were poor in gene density, low in GC content, low in basal expression (except group 0-0.1), and high in H1.2/H1X ratio (**Figure 6D** and **Suppl. Figure 4B-F**). Moreover, the selected TADs were enriched in H1.2 and were poor in H1.4 (**Suppl. Figure 4G**). Interestingly, these TADs suffered less prominent changes in H1 variant distribution or ATAC-seq coverage than non-coordinated response TADs (**Figure 6D,E** and **Suppl. Figure 4H**). Despite this, coordinated TADs were enriched in regions of the genome that suffered decompaction as measured by the Hi-C compartmentalization analysis (**Figure 6F**).

**Figure 6.**
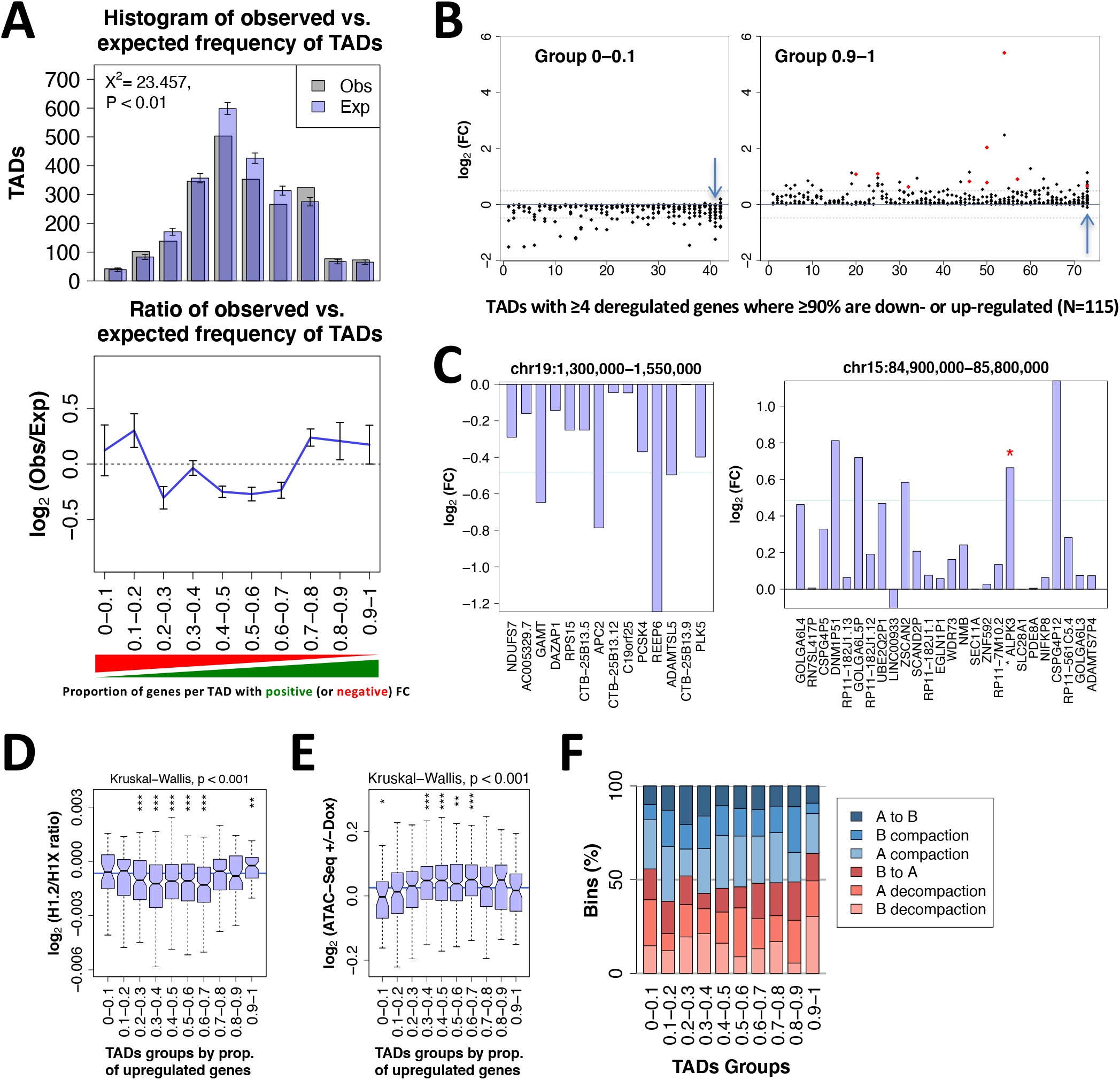
Gene expression is coordinately altered within TADs upon H1 KD. **(A)** *Top-*Histogram of the frequencies of TADs for the observed (gray) or randomized (purple) position of genes, for TADs containing an increasing proportion of genes per TAD with positive FC. Observed and expected values were compared using Pearson’s chi-square test. Gene locations were randomized 10,000 times, constraining by chromosome, not allowing overlapping, and only considering TADs with ≥4 genes. *Bottom-*Ratio of observed versus expected frequencies of TADs with distinct proportions of genes with positive or negative H1 KD-induced FC; FC>1 or FC<-1. **(B)** TADs with ≥4 genes where at least 90% of genes are down-(left) or up-regulated (right) with FC<-1 or FC>1, respectively (total N=115). Log2 of gene expression FC is shown. TADs are ordered from low to high abundance of genes per TAD. Dashed lanes indicate FC=-1.4 or FC=1.4. Red dots represent ISGs. Example genes shown in (C) are located within TADs marked with an arrow. **(C)** Examples of TADs with biased coordinated response to H1 KD. Fold change +Dox/-Dox (log2) is shown for all coding and non-coding genes present within a representative TAD containing 90-100% of genes with negative (left) or positive (right) FC, respectively. Genes are ordered according to their position within the genome. Red asterisk represents ISGs. **(D)** Box plot showing the ChIP-seq H1.2/H1X log2 ratio within TADs in the 10 groups described in (A). **(E)** Box plot showing the ATAC-seq accessibility gain upon H1 KD (+/-Dox) within TADs in the 10 groups described in (A). Kruskal-Wallis test determined that there were statistically significant differences between the groups in (D) and (E). Comparison between each group of TADs and the median ChIP-seq H1.2/H1X log2 ratio (D) or the ATAC-seq accessibility changes (E) was performed using the one-sample Wilcoxon signed-rank test (***) P < 0.001; (**) P < 0.01. **(F)** Bar plots showing the frequency of overlap between all the TAD groups described in (A) and genome segments within A/B compartment categories described in Figure 2C that changed compaction upon H1 KD. The observed and expected count of bins of the different groups of TADs were significantly different (P < 0.001, Pearson’s Chi-squared test).

Altogether, the results support that upon H1 depletion the majority of the genome (53%) does not alter its expression. However, genes located in regions of high H1.2/H1X ratio harbored more genes whose expression was coordinated within entire TADs. Therefore, our results indicate that upon H1 depletion, the entire TADs were architecturally altered and most resident genes were coordinately regulated.

### 3D modeling of TADs with coordinated transcriptional response

To further characterize architecturally changes within TADs with coordinated transcriptional response to H1 KD, we next generated 3D models of genomic regions harboring TADs that contained at least 90% of genes down or up-regulated (group 0-0.1, “*d”*, N=42; group 0.9-1, “*u”*, N=73; **Figure 6B**), both in WT and H1 KD conditions. As a control, we also modeled TADs with the most extreme H1.4 decrease upon H1 KD (group “*h1”*, N=100), TADs with a bidirectional transcriptional response (“*bi”*, N=174, picked from groups 0.4-0.6; **Suppl. Figure 4C**), TADs with minimum gene expression changes (“*mi”*, N=100), and TADs with no annotated genes (“*wi”*, N=12). Models were built based on our Hi-C data at 10 kb resolution using TADbit as previously described^31^. Several structural measures were computed and compared between groups of modeled TADs, such as: consistency, radius of gyration, accessibility, density, and walking angle (**Figure 7** and **Materials and Methods** for the definition of the structural measures). Additionally, the analyzed TAD groups were characterized in terms of H1 abundance, gene expression changes, GC content and ATAC-seq accessibility for comparison with the structural data (**Suppl. Figure 5**). All modeled TAD groups resulted in highly consistent models, this indicates that the input Hi-C data did not contain many contradictory interactions and that fairly structural similar conformations were obtained from the ensemble of models for all cases. Only TADs harboring no genes resulted in 3D models with lower consistency measures indicating that more different conformations could satisfy the input restraints (**Figure 7A**). Interestingly, TADs with the highest H1.4 decrease upon H1 KD (“*h1*” group) as well as TADs with no genes (“*wi*” group) overall resulted in more different structural properties. Specifically, both *h1* and *wi* TADs are more compact (lower radius of gyration) compared to the rest of the groups (**Figure 7A**). However, *h1* results in the densest DNA (bp per nanometer) models compared to the *wi*, which are the least dense of all. Other groups have similar density values and between these two extremes. Finally, the models indicate that upon H1 KD and across all types of TAD groups there is a significant increase of the walking angle measure indicating a change of stiffness of the chromatin (**Figure 7A**). In general, changes in the structural properties of TADs reflected a tendency to chromatin opening upon H1 KD, such as the significant increase of chromatin walking angle and tendency to increase of the radius of gyration. The observed changes are exemplified in three models from the *wi, h1* and *ud* groups (**Figure 7B**). In general, we observed no significant differences in structural changes upon H1 KD in TADs with no genes (*wi*), while the changes were more evident in the *h1* group and also in the *ud* group independently of the direction of the changes in gene expression. Indeed, although without significance, changes in the TADs with a coordinated transcriptional response to H1 KD (*u, d*) have the same trends, indicating that TADs that were coordinately up-or down-regulated were similarly structurally altered upon H1 KD. Our 3D models indicate that all TADs are altered in a similar way due to H1 KD, with different consequences in gene expression deregulation that might depend on local features or distinct H1 abundance.

**Figure 7.**
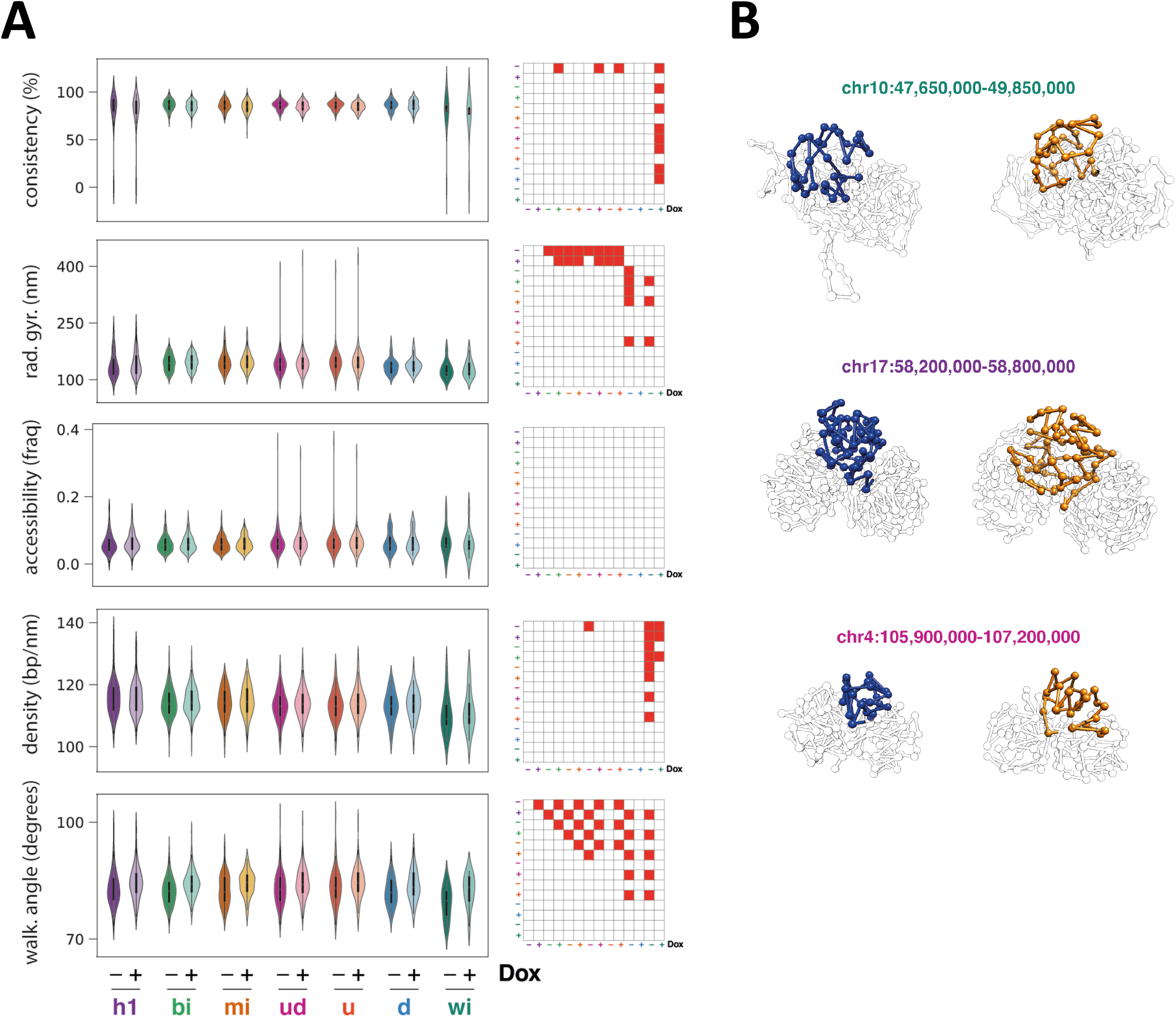
Structural properties of TADs. **(A)** Violin plots of structural properties measured on the 3D models computed for 7 classes of TADs, both in WT and H1 KD conditions (-/+Dox): TADs with the most extreme H1.4 decrease upon H1 KD (*h1*, N=100), TADs presenting a bidirectional transcriptional response to multi-H1 KD (*bi*, N=174), TADs presenting the minimum gene expression changes (*mi*, N=100), TADs presenting a coordinated transcriptional response to multi-H1 KD (*u*, only up-regulated genes, N=73; *d*, only down-regulated genes, N=42; *ud*, TADs *u* and *d* together, N=115), TADs without genes (*wi*, N=12). For each TAD 1,000 models have been generated and clustered, and plotted measures are relative to the main cluster of models. Reported measures are consistency, radius of gyration, accessibility, density and walking angle. Matrices next to violin plots indicate classes of TADs that are significantly different for each measure. Statistical significance of the difference between distributions was computed with Kolmogorov-Smirnov test (p-value<0.01). See Materials and Methods for details. **(B)** 3D models of the indicated TADs within chr10, chr17 and chr4, from the *wi, h1* and *ud* groups, respectively, in WT (blue) and KD (orange) conditions. The 3D modelling reflected a tendency to chromatin opening upon H1 KD.

## DISCUSSION

In this study, we have analyzed the genomic distribution of five endogenous H1 variants within T47D breast cancer cells by ChIP-seq using specific antibodies. This is almost the whole somatic H1 complement of this cell line with the exception of H1.1, which is not expressed in these cells, and H1.3 that was not profiled due to the lack of ChIP-grade antibodies. This is, to our knowledge, the first time that most of endogenous variants have been profiled in a mammalian cell.

In previous studies, we mapped endogenous H1.2 and H1X, demonstrating that they have different distributions across the genome^16,17,20^. On the one side, H1.2 is enriched within intergenic, low gene expression regions and lamina-associated domains. On the other side, H1X is enriched at gene-rich chromosomes, RNA polymerase II enriched sites, coding regions and hypomethylated CpG islands. The apparent differential distribution of the two H1 variants in active versus inactive chromatin, also correlates with the CG content of the regions where they localize. Indeed, we have observed here that H1.5 and H1.0 colocalize with H1.2, at low GC regions, while H1.4 distribution is similar to H1X with the exception of H1X being highly enriched at high GC regions. Previously, we profiled H1.0 and H1.4 fused to an HA tag at C-termini, stably expressed through a lentiviral vector into T47D cells. Using this technique, both H1.4-HA and H1.0-HA were enriched at high GC regions, indicating that profiling exogenous, tagged H1 proteins may give different results than endogenous proteins^16^. In apparent contradiction, H1.0 has been profiled in human skin fibroblasts, being enriched at high GC regions^18^ while in mouse, tagged, knocked-in H1c (H1.2), H1d (H1.3) and H1.0 have been profiled in ESCs and found enriched at low GC regions^14^. Altogether, this suggests that H1 variants distribution might be different among cell types, and could be explained by the relative levels of expression of the different variants. Extensive profiling of H1 variants among different cell types with the same methodology should be done to clarify whether the observed distribution of H1 variants is cell type-specific or universal for some of the variants.

In T47D cells, the H1 content was estimated to be 9% for H1.0, 23% for H1.2, 13% for H1.3, 24% for H1.4 and 31% for H1.5^19^. Our distribution analysis thus indicates that most of H1 variants we profiled are located in low GC regions, which supports its role as heterochromatic protein. However, and as previously described^17^, H1X is enriched at high GC regions suggesting its possible role as regulatory H1. We also found that the enrichment of H1.4 at high GC regions is intriguing as it was suggested that, because of its K26 residue which may be methylated and bind HP1, it could be related to heterochromatin^44,45^. Still, a fraction of H1.4 is at low GC regions, and even at high GC bands it could have a role in repression at particular sites. In fact, when profiled within chromatin states, H1.4 overlapped H3K9me3 distribution, a *bona fide* heterochromatin marker.

To study whether alteration of the total H1 content and relative abundance of the different variants affected the genomic localization of remaining histones, we performed ChIP-seq in T47D cells knocked-down for H1.2 and H1.4 with an inducible system, previously characterized^22^. Interestingly, upon H1 KD, H1.4 preferentially remained at low GC regions, supporting its putative role in heterochromatin, and was displaced from high GC regions. In parallel, H1.5 and specially H1X redistributed to high GC regions. H1.0 maintained its distribution across the genome despite its expression and protein levels increased to compensate the H1 overall ≈30% decrease. Overall, H1.2 was depleted but did not change much its relative genomic distribution. Profiling within chromatin states showed that H1.4 slightly switched towards the H1.2 group upon H1 KD, and H1X decreased at heterochromatin and increased in almost all other chromatin states.

In this work, we have shown that H1 KD caused changes in chromatin accessibility and H3K9me3 distribution, shifts in A/B compartments and TAD borders, and changes in the 3D architecture of TADs (**Figure 8**). Some of these changes were dependent on the compaction or GC content of genomic domains. In fact, we have previously shown that A and B compartments positively correlate with the measured H1.2/H1X ratio^20^. Here we have further shown that the A/B compartments present different abundance of H1 variants and respond differently to H1 depletion. Upon H1 KD, ATAC-seq chromatin accessibility increased genome-wide but more markedly at A compartment. Accordingly, the repressive histone mark H3K9me3 decreased majorly from A compartment. Recent reports have shown that H1 depletion in mouse T cells and germinal centre B cells lead to B-to-A compartment shifting^46,47^. These could be due to the fact that differentiated cells present a well-constituted heterochromatin rich in histone H1, compared to pluripotent and cancer cells where chromatin may be more plastic, partially because of a lower H1 content^48,49^. H1-mediated compartmentalization may be established along differentiation, sequestering the stem cell programs within the B compartment. Deregulation of H1 levels and compartmentalization may occur in cancer and along reprogramming^41,50,51^. The observation of A-to-B and B-to-A shifting in our cancer model T47D cells in similar proportions could be due to an overall less compacted chromatin, or to the simultaneous depletion of H1 variants assayed here to occupy distinct genomic compartments. In mouse cells, so far only tagged H1c (H1.2) and H1d (H1.3) have been ChIP-seq profiled, showing an overlapping distribution within low GC regions/B compartment^14,20^. We here show that H1.2 is abundant at the B compartment and its depletion in multiH1 KD cells resulted in decompaction and B-to-A shifts, accompanied by gene induction and local increase of DNA interactions. Conversely, H1.4 is abundant at the A compartment and its depletion preferentially upon H1 KD accompanied A-to-B shifts or compaction. However, as A decompaction also occurred in regions with H1.4 occupancy, our results could suggest a dual role of this H1 variant, which requires further investigation.

**Figure 8.**
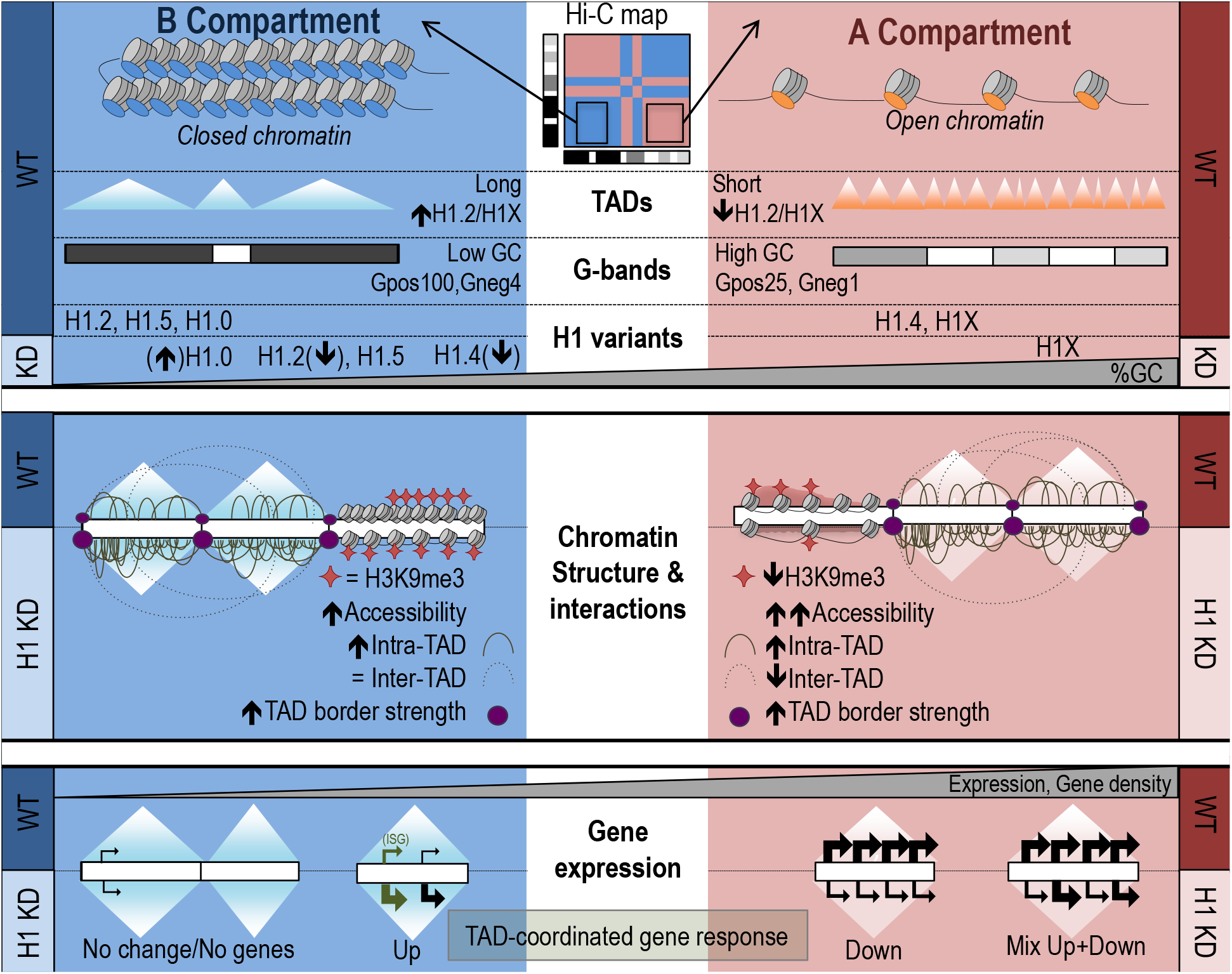
Chromatin organization and consequences upon H1 depletion on genome structure. *Chromatin organization and H1 variants distribution (upper panel):* Hi-C data allows determination of B (inactive) and A (active) compartments. B compartment is characterized by closed chromatin, long TADs with a high H1.2/H1X ratio and a great overlap with low GC Giemsa bands, while the opposite occurs for A compartment. H1 variants were differentially distributed along the genome and two profiles could be distinguished in T47D breast cancer cells: H1.2, H1.5 and H1.0 co-localized at low GC regions whereas H1.4 and H1X occupied high GC regions. Upon multiple H1 depletion (H1 KD), H1.2 and H1.4 were strongly depleted while H1.0 became up-regulated but without changing its distribution. Remaining H1.4 redistributed to low GC regions, whereas H1.2, H1.5 and especially H1X were redistributed to higher GC regions. *Consequences of H1 KD in chromatin structure (middle panel):* Upon H1 KD, chromatin accessibility increased and H3K9me3 signal decreased, especially at A compartment. Intra-TAD interactions increased both at B and A compartments whereas inter-TAD interactions were reduced at A compartment. TAD-border strength increased, together with some TAD borders being lost or shifted. Upon H1 KD, shifts between and within A/B compartments occurred, being more frequent compaction shifts at A compartment (including A-to-B shifts) and decompaction at B compartment (incl. B-to-A). *Consequences of H1 KD in genome structure are related to gene expression deregulation (bottom panel):* TADs presenting a coordinated response to H1 KD were enriched compared to the expected frequency. Up-regulated genes accumulated within TADs with poor basal expression and low gene density. Gene-dense TADs contained both up- and down-regulated genes simultaneously. TADs with only down-regulated genes showed intermediate features.

Previously, we and others have shown that epigenetic states and H1 distribution are more homogeneous within a TAD, suggesting that TAD borders prevent the spreading of these features^6,20^. In our work, TADs hardly changed its size or distribution upon multiH1 KD, however, a clear increased TAD border strength and intra-TAD contacts was observed. Interestingly, the concomitant inter-TAD contacts reduced more predominantly in A compartment compared to the B compartment. Indeed, several reports have also shown that TAD organization remains largely unchanged when disturbing chromatin homeostasis, including mouse H1-depleted cells or epithelial-to-mesenchymal transition^6,30,52,53^. However, our work now highlights novel relevant changes in TAD organization due to depletion of H1 variants, including an increase in border strength accompanied by an increase of intra-TAD interactions.

We have shown here and previously that H1 variants selective depletion results in changes in expression of hundreds of genes, including de-repression of intergenic and intronic RNAs, as well as heterochromatic repeats and ERVs, which leads to the induction of the interferon response^22^. Moreover, we have shown that responsive genes are non-randomly located throughout the genome but enriched in a limited number of TADs with their resident genes coordinately changing expression to H1 depletion. We have previously reported that, upon H1 KD in T47D cells, the interferon response is induced with many ISGs being up-regulated. This is due to the accumulation of RNAs from repeats and ERVs, which stimulate the response at cytoplasm mimicking a viral infection and resulting in the transcription of many genes involved in such response. A part of this direct effect of H1 depletion, our results may also indicate that other genes not directly related to such response may be deregulated due to structural changes, chromatin decompaction, or simply by co-existing within the same TAD with genes directly activated. Indeed, we show that ISGs up-regulated genes co-exist within TADs with other genes that coordinated respond to H1 KD. Despite this observation, we also found that many TADs with a coordinated response do not contain annotated ISGs genes, so we propose that the response may be a consequence of architectural changes upon H1 KD. This result is further supported by the 3D modeling of TADs.

Overall, our results indicate that the non-random genomic distribution of H1 variants, their re-location upon variant depletion, and the subsequent genome structural changes have a read-out in their direct (but also indirect) change of the gene expression program.

## DATA AVAILABILITY STATEMENT

Hi-C and ChIP-seq data on T47D breast cancer cells reported here and deposited in NCBI’s Gene Expression Omnibus are accessible through GEO Series accession number GSE172618 and GSE156036/GSE166645, respectively.

## SUPPLEMENTARY DATA

Supplementary Data are available online.

## ACKNOWLEDGEMENTS

We acknowledge Generalitat de Catalunya and the European Social Fund for AGAUR-FI predoctoral fellowships [to M.S.-P. and to F.M.].

