## Supplemental Table1 and Figures 1 to 5 for "Coordinated changes in gene expression, H1 variant distribution and genome 3D conformation in response to H1 depletion"

#### SUPPLEMENTARY MATERIAL

**Supplementary Table 1.** Hi-C experimental statistics. Statistics shown separated by replicates (top table) and merged datasets (middle table for valid pairs and bottom table for filtered reads).

#### SUPPLEMENTARY FIGURE LEGENDS

**Supplementary Figure 1. Characterization of ChIP-grade H1 antibodies and multiH1 KD cells.** The specificity of H1 antibodies not used before was further tested on Western blot and ChIP-qPCR with chromatin extracts from HeLa and HCT-116 cells, which lack H1.0 or H1.5, respectively, and with chromatin extracts from H1.0, H1.4 or H1.5 single KD T47D cells**. (A)** Immunoblot analysis of H1 variants expression within cell lines. Chromatin extracts (10 µg of protein) from T47D, HeLa and HCT-116 cells were run in SDS/PAGE and immunoblotted with the indicated antibodies against histone H1 variants or histone H3 as a loading control. **(B)** ChIP-qPCR of H1 variants in HeLa and HCT-116 cells. Chromatin was used for ChIP with antibodies against H1 variants and unrelated IgG as a control. Resulting DNA was amplified by qPCR with oligos for distal promoter (3kb upstream TSS) and TSS regions of genes CDK2 and NANOG. ChIP amplification is shown relative to input DNA amplification. A representative experiment is shown. **(C)** Immunoblot analysis of H1 variants expression in inducible knock-down T47D cells for different H1 variants. Chromatin extracted from untreated or 6-days-Dox-treated single-H1 (H1.0, H1.4 or H1.5) KDs cells was immunoblotted with the indicated antibodies or Coomassie stained as a loading control. T47D single-H1 KDs performance was previously characterized in Sancho et al., 2008 (ref.19 of manuscript). **(D)** ChIP-qPCR of H1 variants in inducible knock-down T47D cells for different H1 variants. Chromatin samples shown in (C) were used to perform ChIP with antibodies against H1 variants. Resulting DNA was amplified by qPCR with oligos for distal promoter (3kb upstream TSS) and TSS regions of genes CDK2 and NANOG. ChIP amplification is shown relative to input DNA amplification. A representative experiment is shown. **(E)** Further, the effect of H1 KD was confirmed by RT-qPCR, showing inhibition of multiple H1 variant genes and induction of satellites, repeats and interferon stimulated genes (ISGs). RNA extracted from T47D multiH1 KD cells cultured in the presence or not of Doxycycline for 6 days was reverse transcribed with random hexamers. Resulting cDNA was submitted to real-time PCR with oligos to measure expression of H1 variants genes, ISGs (IFI27 and OASL), satellites (SATa) and endogenous retroviruses (MER4D, MER21C and MLT1C49). Expression was represented relative to GAPDH as a control and relative to untreated cells. A representative experiment quantified in triplicate is shown.

**Supplementary Figure 2. Genomic distribution of five histone H1 variants in breast cancer cells. (A)** Box plots showing H1 variants input-subtracted ChIP-seq abundance within G bands, for each band type (Gneg1-4 and Gpos25-100), in multiH1 KD cells treated or not with Doxycycline. GC content of G bands for each band type is also represented. Gneg bands were divided in 4 equal groups according to GC content. Wilcoxon signed-rank test was used to compare the H1 variants abundance within G bands before and after multiH1 KD: (***) $P < 0.001$; (**) $P < 0.01$; (*) $P < 0.05$. Gpos25, N = 87; Gpos50, N = 121; Gpos75, N = 89; Gpos100, N = 81; Gneg, N = 414. Data corresponds to a representative ChIP-seq experiment. **(B)** Scatter plots of the indicated H1 variant pairs input-subtracted ChIP-seq abundance within 100-kb bins of the human genome. The GC content at each bin is color-coded. Pearson's correlation coefficient is shown (*P*-value<0.001). **(C)** Scatter plot of H1.2 and H1X input-subtracted ChIP-seq abundance within Gpos and Gneg bands from T47D multiH1sh untreated cells. Pearson's correlation coefficient is shown (*P*-value<0.001). **(D)** Heat map and cluster analysis of the input-subtracted ChIP-seq abundance of H1 variants from WT or H1 KD T47D cells (-/+Dox) within a random sample of genome fragments belonging to the 10 chromatin states. For each chromatin state 1,000 fragments were randomly picked and 50 groups of 20 fragments were randomly generated. Each lane of the heat map represents the median input-subtracted ChIP-seq abundance of H1 variants in a group. Clustering confirmed that H1X was the most extreme variant and H1.4 showed an intermediate distribution together with H3K9me3.

**Supplementary Figure 3. Hi-C interactions map of H1 KD T47D cells.** Hi-C interaction maps for all 3 replicates and the merged maps of WT and H1 KD. Left plots correspond to WT maps and right plots are H1 KD maps. Rows show maps for replicates 1 to 3 as well as the merged maps. Each panel includes a Hi-C raw interaction maps at 1Mb resolution and 100 kb resolution for genome-wide and chromosome 1, respectively.

**Supplementary Figure 4. Characterization of TADs presenting a coordinated regulation of gene expression upon H1 KD. (A)** TADs with ≥4 genes where at least 80% of genes are down- (left) or up-regulated (right) with FC<-1 or FC>1, respectively, upon H1 KD (total N=294). Log2 of gene expression fold-change (FC) is shown. TADs are ordered from low to high abundance of genes per TAD. Dashed lanes indicate FC=-1.4 or FC=1.4. Red dots represent ISGs. **(B)** TAD groups by proportion of genes per TAD with positive FC upon H1 KD. Only TADs with ≥4 genes were considered. **(C-D)** Box plots showing the expression levels -/+Dox (C) and fold-change (D) within TADs in the 10 groups described in (B). The number of TADs within each group is indicated in (C). (***) $P < 0.001$ (Wilcoxon signed-rank test). **(E-F)** Box plots showing the GC content (E) and number of genes per TAD (normalized to TAD length and multiplied by the average TADs length) (F) in the 10 TAD groups described in (B). **(G)** Box plots showing the H1.2 and H1.4 ChIP-seq signal in untreated cells within TADs in the 10 groups described in (B). **(H)** Box plots showing the H1 variants input-subtracted ChIP-seq signal ratio in H1 KD (+Dox) compared to WT (-Dox) cells within TADs in the 10 groups described in (B). Kruskal-Wallis test was applied in all boxplot analysis (D-H) to determine if there were statistically significant differences between the groups. One-sample Wilcoxon signed-rank test was subsequently applied to compare each group of TADs against the median value for each analyzed property.

**Supplementary Figure 5. Properties of TADs structurally analysed.** Properties of the 7 classes of TADs described in Figure 7 in WT (-Dox) and H1 KD (+Dox) conditions: GC content, expression of genes within TADs, input-subtracted H1.2 and H1.4 ChIP-seq signal, ATAC-seq signal or normalized peaks count; as well as changes +Dox/-Dox within the TAD groups (log2 ratio) on gene expression, H1.4 signal, and ATAC-seq signal. The differences between WT and H1 KD conditions were examined using the Wilcoxon signed-rank test. The global differences between the groups of TADs showing +Dox/-Dox changes were determined by applying a Kruskal-Wallis test. One-sample Wilcoxon signed-rank test was subsequently applied to compare each group of TADs against the median value for each analyzed property. (***) $P < 0.001$; (**) $P < 0.01$; (*) $P < 0.05$.

**Supplementary Table 1. Hi-C experimental statistics. Statistics shown separated by replicates (top table) and merged datasets (middle table for valid pairs and bottom table for filtered reads).**

| Sample | Total reads | Mapped read-end 1 | Mapped read-end 2 | Mapped reads | % mapped | Valid reads | % valid |
|---|---|---|---|---|---|---|---|
| Rep1 -Dox | 342,288,245 | 259,075,154 | 258,107,758 | 208,117,630 | 60.8 | 152,825,989 | 44.65 |
| Rep2 -Dox | 174,269,331 | 135,986,461 | 135,602,178 | 109,694,504 | 62.95 | 93,012,379 | 53.37 |
| Rep3 -Dox | 151,794,765 | 117,907,548 | 117,082,420 | 95,250,499 | 62.75 | 75,565,442 | 49.78 |
| Rep1 +Dox | 326,519,453 | 250,463,260 | 248,485,925 | 201,425,511 | 61.69 | 155,955,437 | 47.76 |
| Rep2 +Dox | 189,256,710 | 148,401,462 | 147,801,442 | 120,043,103 | 63.43 | 102,262,949 | 54.03 |
| Rep3 +Dox | 149,483,281 | 117,243,624 | 116,902,071 | 95,143,251 | 63.65 | 80,454,962 | 53.82 |

**Separated replicates**

**Merged replicates – valid reads**

| Sample | Total reads | Valid reads | % valid |
|---|---|---|---|
| Rep123 -Dox | 668,352,341 | 321,403,810 | 48.09 |
| Rep123 +Dox | 665,259,444 | 338,673,348 | 50.91 |

**Merged replicates – filtered artifacts**

| Sample | Random breaks | Self-circle | Too close from RES | Over-represented | Dangling-end | Too large | Extra dangling-end | Error | Too short | Duplicated |
|---|---|---|---|---|---|---|---|---|---|---|
| Rep123 -Dox | 3,057,406 | 399,186 | 99,467,116 | 15,284,567 | 41,896,379 | 319,233 | 32,839,003 | 1,344,033 | 11,543,359 | 7,539,158 |
| Rep123 +Dox | 2,750,619 | 394,632 | 94,255,761 | 13,603,507 | 33,351,995 | 363,675 | 28,989,911 | 166,760 | 11,403,959 | 6,818,581 |

### Suppl. Figure 1

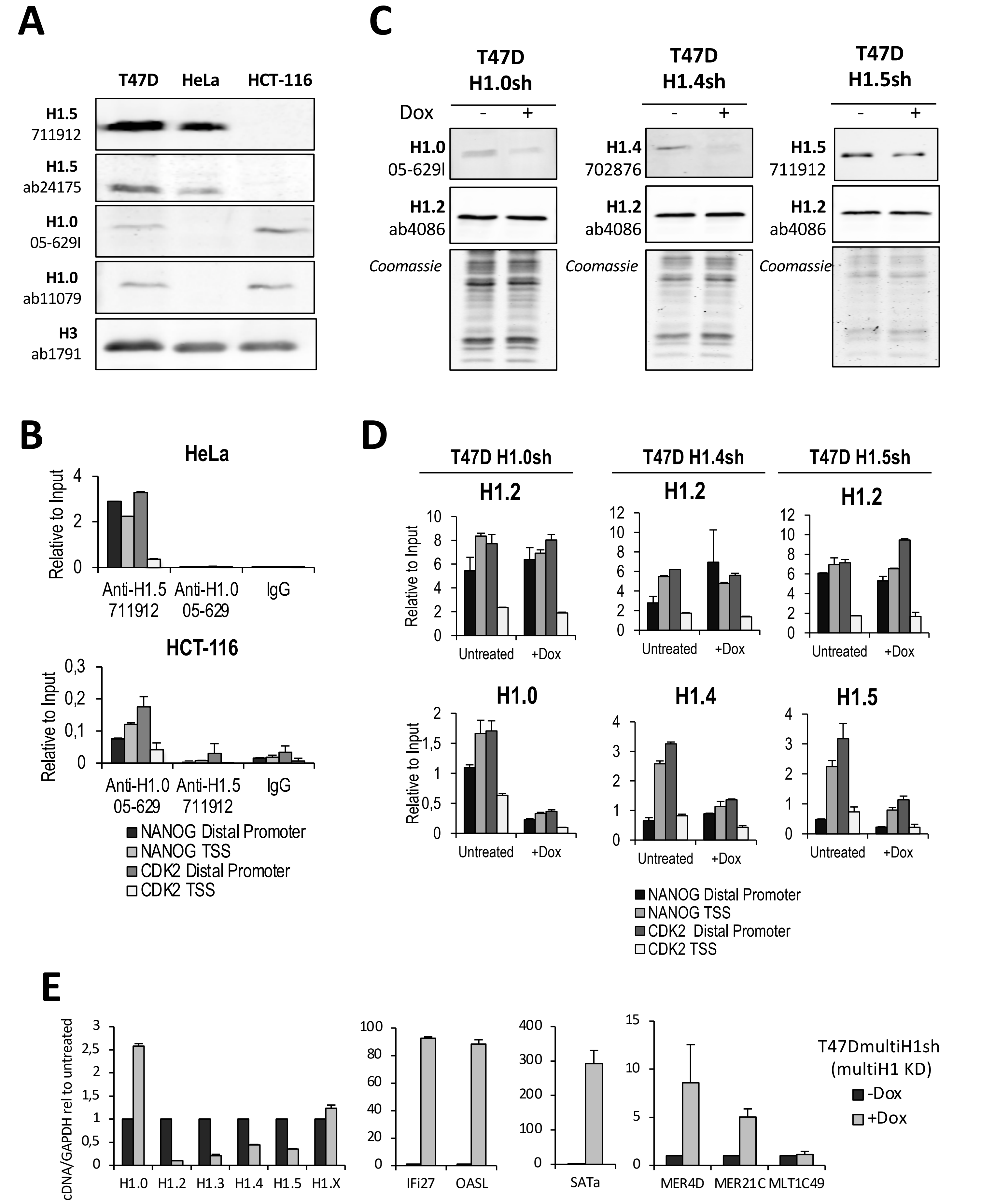

### Suppl. Figure 2

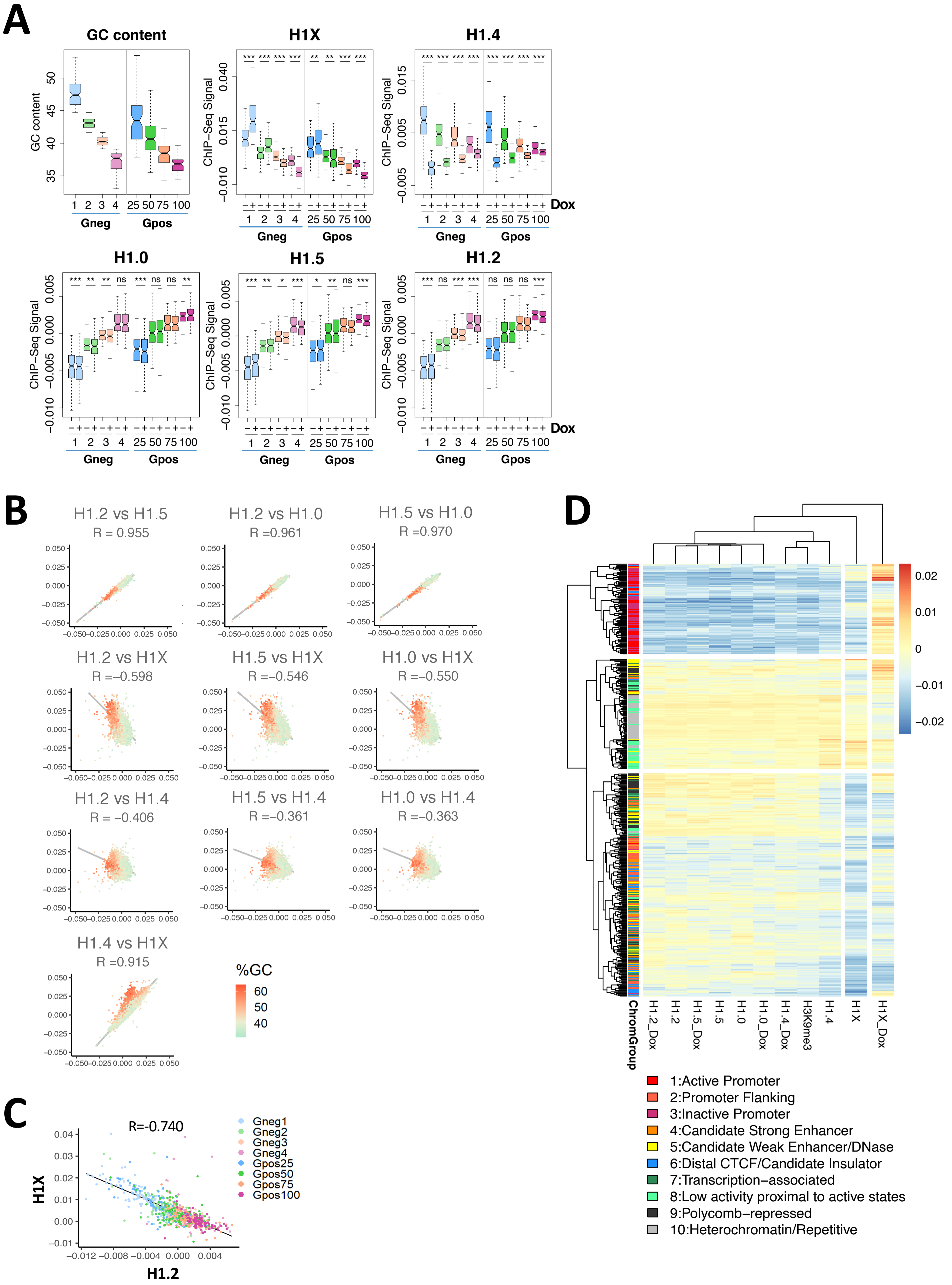

### Suppl. Figure 3

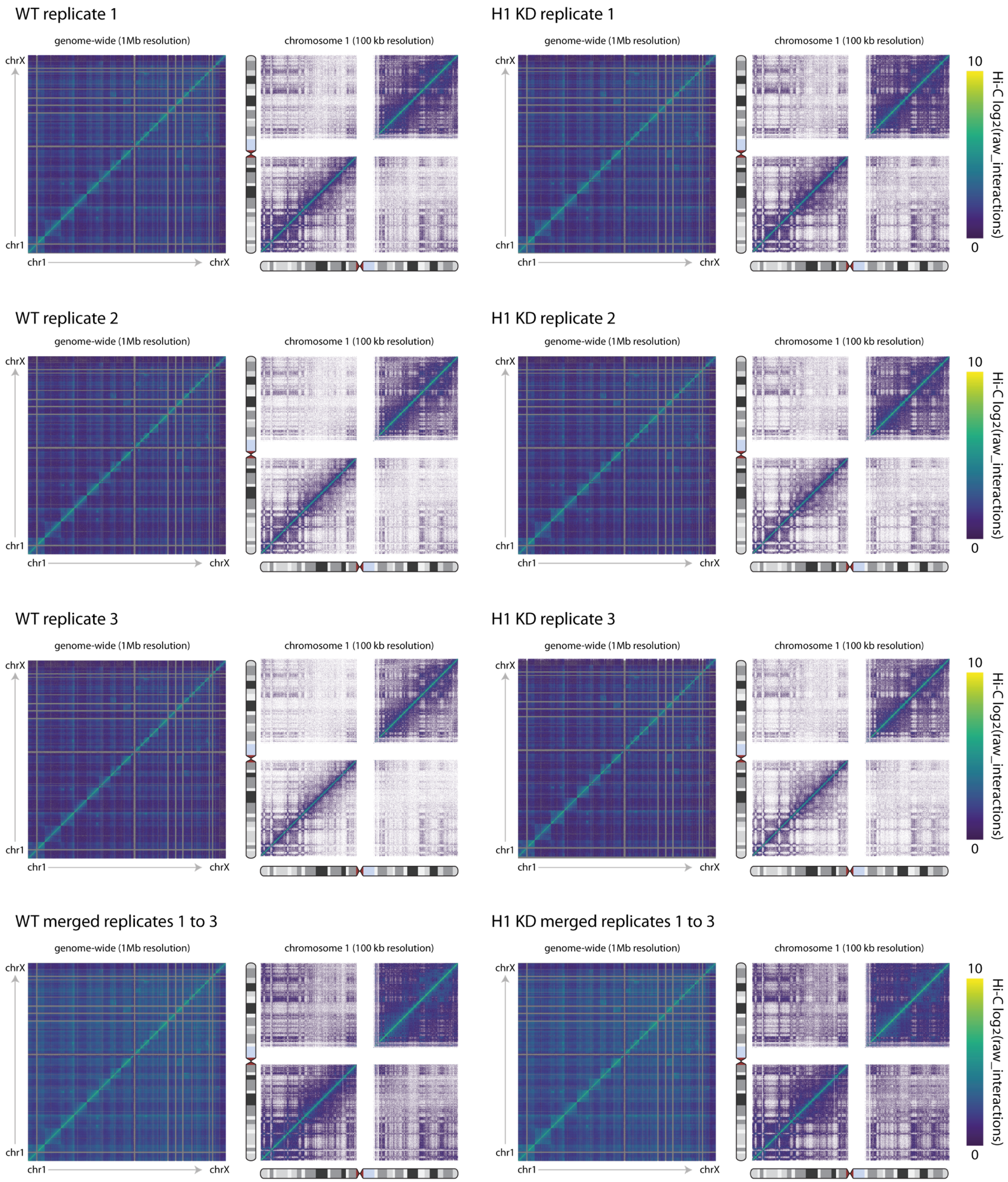

### Suppl. Figure 4

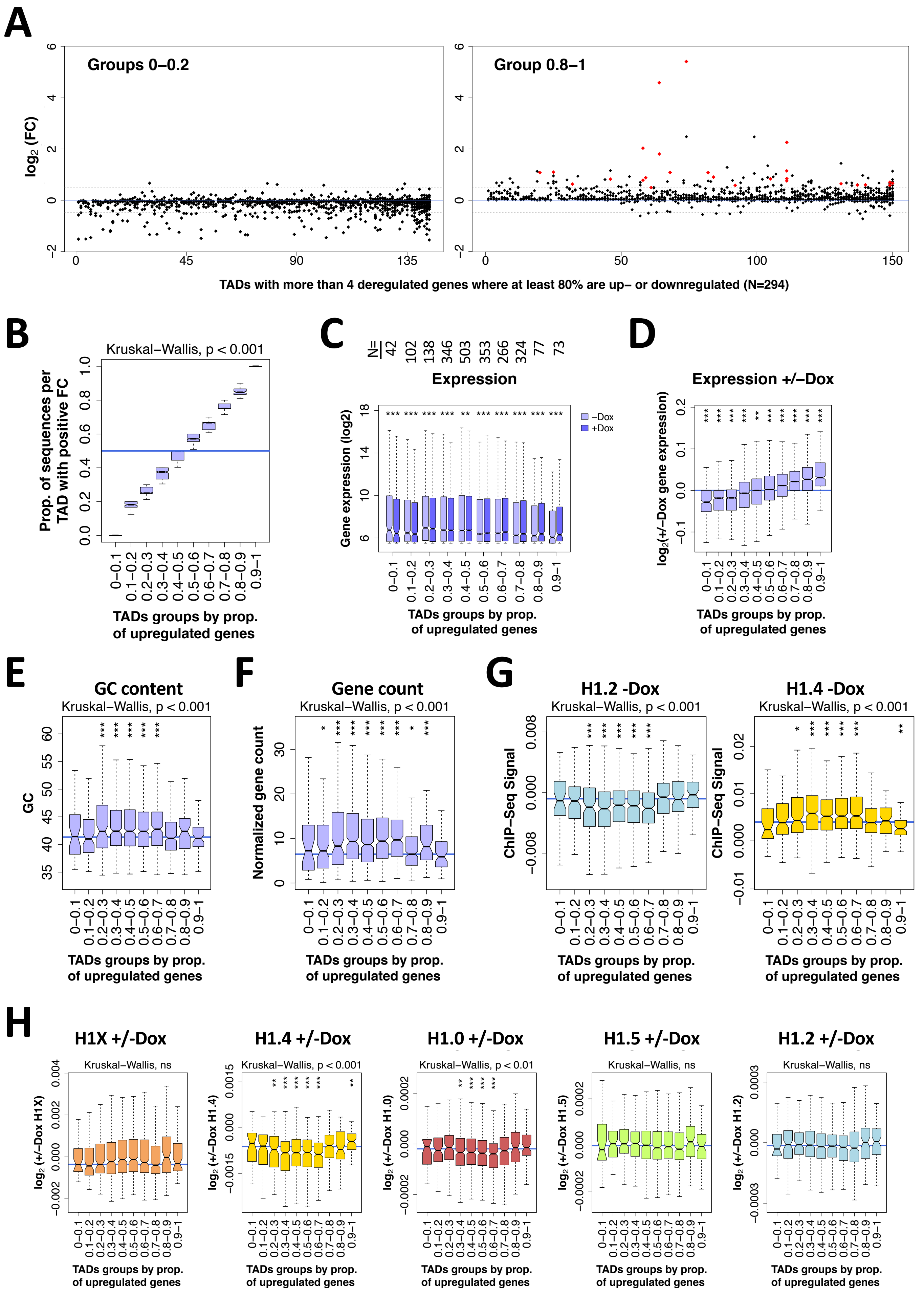

### Suppl. Figure 5

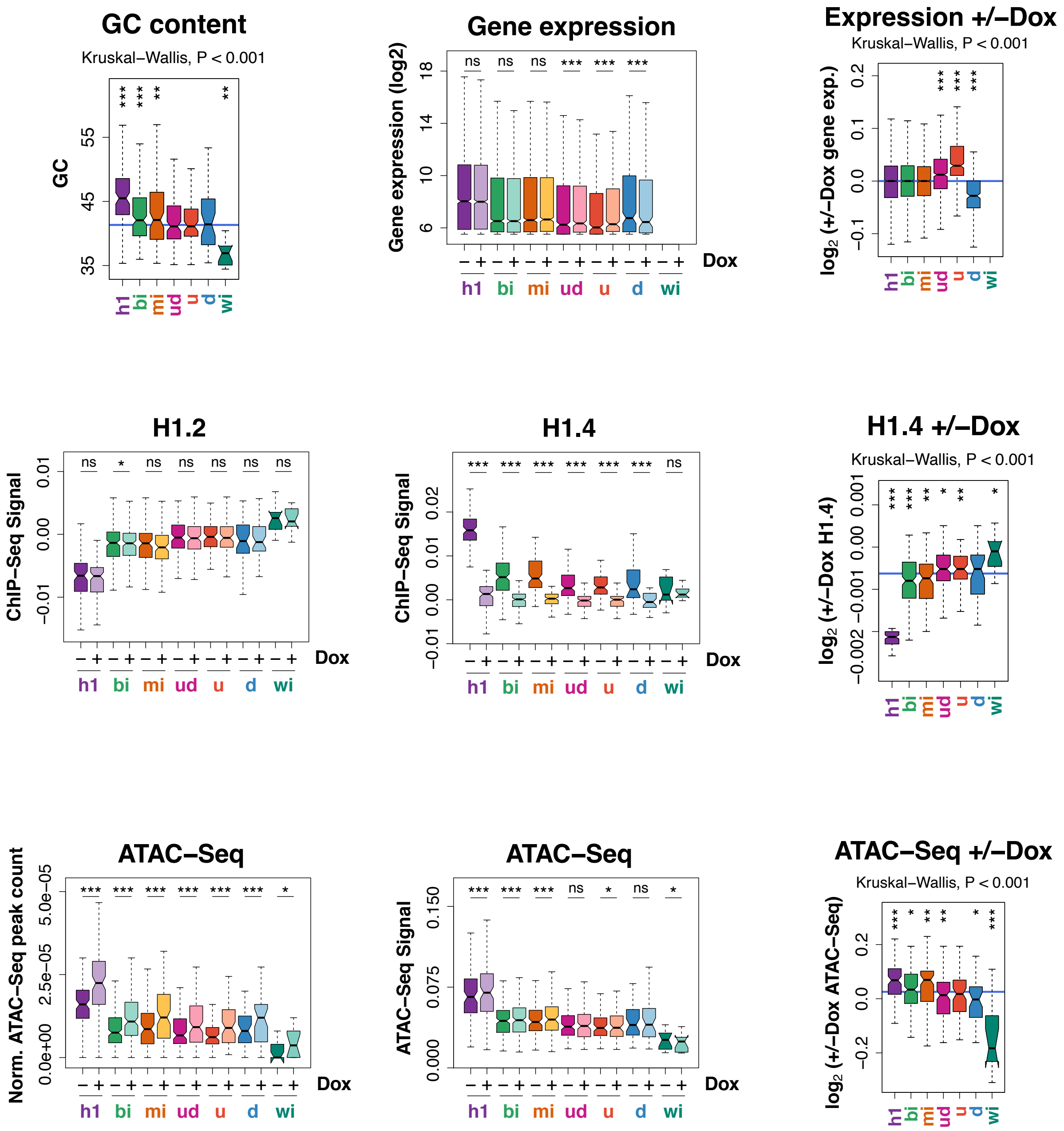
